# Human mascRNA interacts with SUMO E1 and reshapes nuclear SUMOylation

**DOI:** 10.64898/2026.01.19.700226

**Authors:** Eleni Liapi, Kaili Dawn Taylor, Aleah Marissa Holts, Joëlle Rudinger-Thirion, Claude Sauter, Anastasios Vourekas

## Abstract

Human mascRNA is a tRNA-like 58-nucleotide non-coding RNA processed from the 3′ end of the MALAT1 transcript. Although mascRNA has been implicated in diverse cellular processes, its molecular mechanism has remained unclear. Here, we reveal a direct functional link between mascRNA and the SUMOylation pathway. Structural probing and SEC-SAXS confirm the compact, tRNA-like architecture of mascRNA in solution, while biochemical assays demonstrate its direct interaction with the SUMO E1 activating enzyme *in vitro*. Proteomic analyses show that mascRNA engages nuclear protein networks enriched for SUMOylated factors, including chromatin remodelers and nucleolar components. Transfection of synthetic mascRNA alters SAE2 and SUMO1 nuclear organization and broadly reduces nuclear SUMOylation, accompanied by decreased ProMyeLocytic (PML) body abundance and SUMO colocalization. SUMO1 immunoprecipitation and mass spectrometry reveal widespread decreases in SUMO conjugation, particularly among proteins involved in RNA metabolism and rRNA biogenesis with nuclear body localization. We further validate reduced SUMOylation of nucleolar factors such as BEND3 and ZNF106. These findings position mascRNA as a structured RNA ligand that modulates SUMO-dependent nuclear architecture and protein homeostasis, uncovering an unexpected intersection between RNA biology and post-translational modification networks.

**GRAPHICAL ABSTRACT:** 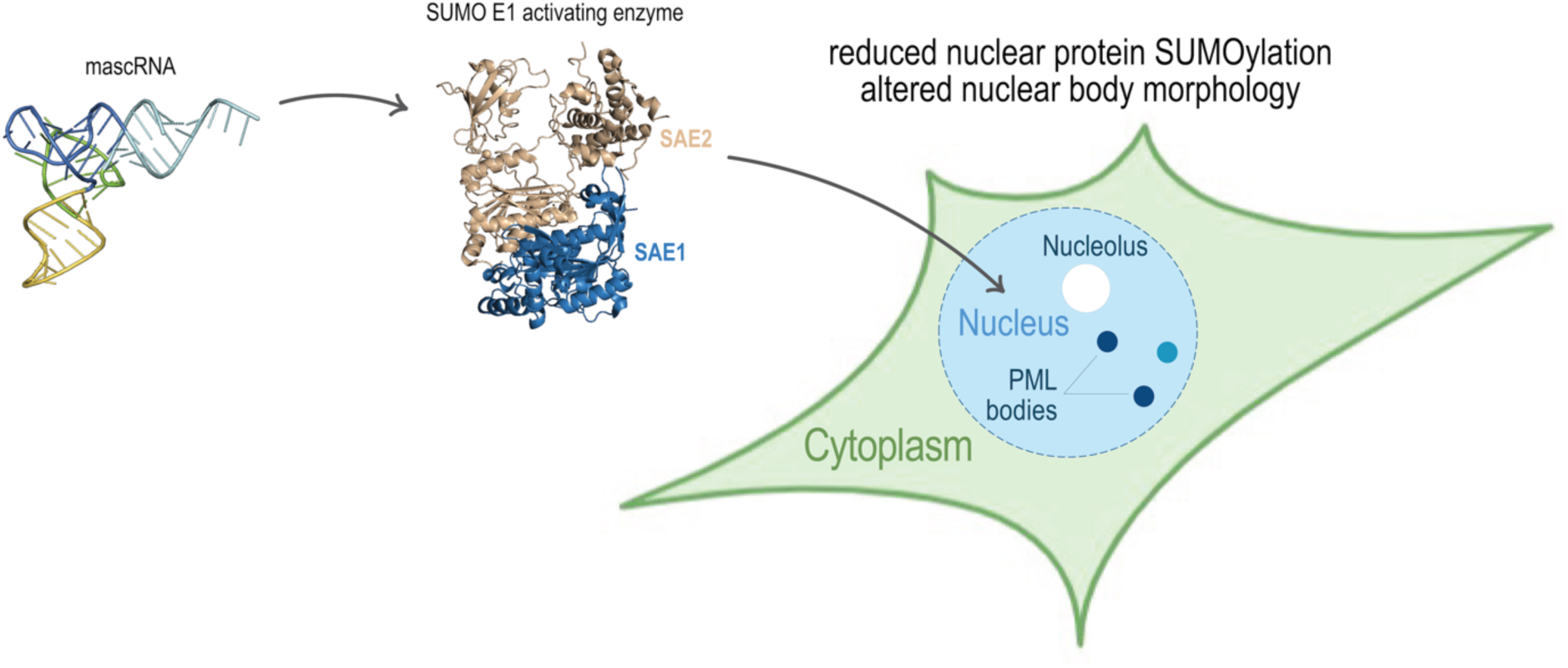

## INTRODUCTION

Human MALAT1-associated small cytoplasmic RNA or mascRNA is a 58 nt non-coding RNA produced via RNase P directed cleavage from the 3’ end of MALAT1 (Metastasis Associated Lung Adenocarcinoma Transcript 1) long <u>n</u>on-coding <u>R</u>NA (lncRNA) primary transcript, followed by RNase Z cleavage of the 3’ trailer sequence, 3’ CCA addition, but no aminoacylation (1). Early 2D predictions (1), SHAPE/DMS data (2) and crystal structures (3, 4) support that mascRNA adopts a tRNA fold, and antibody-based mapping showed that it is chemically modified (methylated) at A43 (5). Moreover, an interaction of the 3’ trailer region downstream of mascRNA is necessary for RNase P to act on nascent MALAT1 transcript and to drive proper formation of MALAT1 3’ triple helix (6). Thus, processing of the MALAT1 primary transcript yields mature MALAT1 and mascRNA as distinct RNA species with different functions and subcellular distributions: MALAT1 remains nuclear and localizes mainly in the nuclear speckles (7), while mascRNA has been reported to be mainly cytoplasmic (1, 8).

mascRNA has been implicated in diverse biological processes, including cell cycle progression in the context of cancer (9), viral replication (10), immune responses (11) and upregulation of global protein translation (8). Exogenously administered mascRNA prevented the replication of the myocarditis-inducing CVB3 RNA virus in both murine and rat cardiomyocytes *ex*-vivo (10). The same virus, CVB3, was reported to induce SUMOylation of endogenous protein REGγ (a proteasome activator) by impacting its translocation to cytoplasm, which subsequently led to p53 degradation and relief of the restriction for viral replication (12). Interestingly, CVB3 infection increased levels of SUMO peptidase Senp2, a finding that implicates the SUMO protein machinery in the mammalian host response towards this viral challenge (13). CVB3 levels were higher in cardiomyocyte-specific *Senp2* KO than WT mice (13), suggesting that the balanced/coordinated function of the SUMO PTM is essential for limiting CVB3-infection. Recently, Goffeney *et* al., demonstrated a molecular circuitry on how SUMOylated MORC3 controls IFNB1 expression in the context of 3D chromatin reorganization in myeloid cells (14). The SUMOylation post-translational modification (PTM) cascade is heavily involved in immune responses during viral infections resulting in changes in biological condensates and protein dynamics (15, 16). However, since the initial discovery of mascRNA limiting CVB3 viral RNA levels, the molecular mechanism of function mediated by protein nodes remains incompletely understood.

The SUMOylation PTM is critical for nuclear processes, including cell cycle control (17), host antiviral responses, frequently acting in crosstalk with the ubiquitination pathway (18). The SUMO pathway is initiated by the activation step, where a SUMO-modifier peptide is loaded onto the SAE2 subunit of the SUMO E1 activating enzyme (heterodimer SAE1/SAE2) (19), followed by the transfer to the UBC9 (the E2 ligase), with or without involvement of an E3 ligase) and ends with the SUMO-conjugation onto lysine residues of recipient protein substrates (20).

The SUMOylation cascade exhibits crosstalk with ubiquitination, with deubiquitinases being prime substrates for this dual co-regulation (21, 22). Emerging studies have implicated mascRNA in the modulation of STING ubiquitination during DNA virus replication. (23). Hu *et* al. reported that in macrophages, exogenous mascRNA limited lung inflammation induced by a DNA virus, a process attributed to STING protein K48 ubiquitination (23). Similarly, Sun *et* al., showed that mascRNA drove K48-Ubiquitination of TRAF6 protein and modulated the TLR vs TRIF-based immune signalling (11). The same study reported that in cytoplasmic lysates of LPS-stimulated (and thus, differentiated) THP1-macrophages (but not THP-1 monocytes), mascRNA associated with RBPs hnRNP H/F. Deregulated immune signaling was similarly reported in a study by Gast *et* al., 2022, using a partially deleted mascRNA in the genome of human THP-1 monocytes (24). The mechanistic details of these mascRNA functions are not yet elucidated.

Taken together, these findings led us to hypothesize that the SUMOylation pathway is implicated in the cellular function of mascRNA. Given accumulating evidence that the SUMOylation cascade plays a central role in immune responses during viral infection and engages in extensive crosstalk with ubiquitination, we further posited that mascRNA functionally associates with components of the SUMOylation machinery, enzymes and/or substrates, and modulates this post-translational modification pathway.

Here, we integrate RNA structural, biochemical, and proteomic analyses to define a functional link between mascRNA and the SUMOylation pathway. We show that mascRNA adopts a tRNA-like cloverleaf and an L-shaped tertiary fold in solution and directly associates with the SUMO E1 activating enzyme via SAE2 both *in vitro* and in cells. Exogenous mascRNA alters nuclear SAE2 and SUMO1 organization and engages predominantly nuclear protein networks enriched for SUMOylated factors. Proteomic analysis of SUMO1-associated proteins further reveals that mascRNA broadly dampens nuclear SUMO1 conjugation, accompanied by reduced PML body abundance and SUMO colocalization. Finally, we validate decreased SUMOylation of nucleolar proteins involved in rRNA biogenesis, supporting a model in which mascRNA modulates SUMO-dependent nuclear body architecture and protein homeostasis.

## MATERIAL AND METHODS

### Radiolabeling of RNA molecules

For autoradiography experiments, 20 pmol of human synthetic mascRNA oligonucleotide without 5’ phosphate group (synthesized by IDT, purified by reverse-phase HPLC), was used as substrate for 5’ end radiolabeling, using the protocol for T4 Poly Nucleotide Kinase (PNK) (NEB M0201S) with 50 pmol [γ-^32^P] ATP (Perkin Elmer BLU002A250UC), 2 μL T4 PNK reaction buffer, 20 units of T4 PNK enzyme (M0201S, NEB), 0.5 μL rRNAsin inhibitor (N2515, Promega) 8.5 μL Nuclease-free water (AM9930, Invitrogen) to a final volume of 20 μL. The radiolabeling reaction was incubated at 37 °C for 30 min. Next, 1 μL of “cold” (non-radiolabeled) 10 mM ATP (Thermo Scientific R0441) was added to phosphorylate any remaining 5’ends, for 5 min at 37 °C. The reaction was heat-inactivated at 65 °C for 20 min and labeled RNAs were retrieved after a G-25 micro-spin column (GE Healthcare Illustra 27532501) step.

### Temperature-folding of RNA molecules

MascRNA and other oligonucleotides were allowed to refold in solution after brief thermal denaturation, before enzymatic foot-printing or other biochemical assays, such as EMSA. Temperature-refolded and unfolded mascRNA were compared to infer accessibility and adoption of tRNA-like structure. The 5’ radiolabeled RNA molecules were refolded in the absence of Mg^2+^ as follows: 1 min at 95 °C with 5 °C drop/15 sec until 25 °C, and then kept on ice.

### Enzymatic probing of mascRNA structure

The structural mapping of synthetic mascRNA transcript was performed using both enzymatic (RNases S1, T1, and V1) and chemical (lead acetate) probes. The radiolabeled mascRNA (4 µg) was purified by electrophoresis on a 12% denaturing polyacrylamide gel, recovered by gel elution and resuspended in water to obtain 50000 cpm /µl. We observed that the reverse-phase HPLC purified mascRNA oligo migrated as a single band but more slowly than predicted on denaturing urea-PAGE (∼70-nt apparent size), consistent with anomalous electrophoretic mobility observed for structured RNAs. Nuclease digestions were carried out at 25 °C for 10 minutes in a buffer containing 25 mM KCl, 10 mM MgCl₂, and 50 mM HEPES-KOH pH 7.5. For RNase S1, the buffer was supplemented with 1 mM ZnCl₂. Reaction mixtures (10 µl) contained 50000 cpm of radiolabeled transcript, 10 pmol of unlabeled transcript, and the following nuclease amounts: 0.25 U RNase T1, 7.5 U RNase S1, or 7 × 10⁻⁴ U RNase V1. Reactions were stopped by phenol/chloroform extraction, followed by ethanol precipitation. Lead acetate probing was performed in 10 µl reactions at 25 °C in a buffer containing 50 mM HEPES-KOH (pH 7.5), 5 mM magnesium acetate, and 50 mM potassium acetate. Each reaction included 50000 cpm of radiolabeled mascRNA transcript and 20 pmol of unlabeled transcript. Freshly prepared lead acetate was added to final concentrations of 1, 2.5, 5, or 10 mM. After 8 minutes, reactions were quenched by addition of 30 mM EDTA and 1 µg glycogen. Samples were ethanol precipitated, and the resulting pellets were washed, dried, and resuspended in 10 µl of formamide dye. Prior to electrophoresis, samples were heated at 90 °C for 2 minutes and then loaded onto a 14% m/v denaturing polyacrylamide gel. A no-lead control was included. Cleavage products were assigned by comparison with alkaline hydrolysis and RNase T1 digests under denaturing condition (25).

### SAXS analysis

Small-angle X-ray scattering (SEC-SAXS) experiments were performed on the SWING beamline (26) at Synchrotron SOLEIL (Saint-Aubin, France) with a beam wavelength of λ = 1.033 Å and the EigerX4M detector (Dectris) positioned at a distance of 2, 000 mm from the sample with the direct beam off-centered. The resulting exploitable q-range was 0.005 - 0.55 Å^-1^, where the wave vector q = 4π sin θ / λ and 2θ is the scattering angle. Samples were loaded onto a BioSec3-300 (Agilent Technologies) column installed upstream the SAXS cell. Experiments were performed at 15°C with a column pre-equilibrated with 50 mM HEPES-NaOH pH 7.5, 150 NaCl, 10 mM MgCl2 and SAXS measurements were performed throughout elution operated at a flow rate of 0.3 ml/min. Data were collected with a frame duration of 990 ms and intervals of 10 ms. 180 frames were collected to estimate the buffer contribution (to be subtracted) and 660 frames in the sample elution window. The mascRNA transcript used for SEC-SAXS experiments included the 3′-CCA sequence and was produced according to Fechter et al (27). Purification was performed by denaturing polyacrylamide gel electrophoresis followed by electroelution, as previously described (28). A total of 40 µl of mascRNA solution at 2.8 mg/ml (110 µg) was injected onto the SEC column and scattering signal was collected throughout the elution. Data processing was performed using Foxtrot (29), and analysis using BioXTAS-RAW 2.3 (30) and ATSAS 4.0.1 packages (31). Radius of gyration was estimated with the Guinier analysis. It was also derived, together with the maximal dimension (Dmax) from the distance distribution function p(r) of the particle using the indirect transform package GNOM (32). Particle compactness was assessed from the normalized Kratky plot in BioXTAS-RAW. Models of mascRNA were produced using Assemble2 (33), trRosettaRNA (34), and AlphaFold3 (35). Their theoretical SAXS profile was compared to the experimental SAXS curves owing to a goodness-of-fit calculated with CRYSOL (32). Graphics were prepared with LibreOffice Draw (The Document Foundation) and BioXTAS-RAW.

### Heterologous expression and affinity purification of SAE1/2 complex

Recombinant human 6xHis-SAE1/SAE2 complex (custom order, Genscript) was expressed in *E. coli* BL21 Star™ (DE3) from codon-optimized sequences cloned into the pET28a for SAE1 and pET11d for SAE2 with N-terminal His tags for both proteins. Expression was induced with IPTG in TB medium at 16 °C for 16 h, and cells were harvested by centrifugation. The protein complex was purified from clarified lysates by sequential Ni-affinity, Superdex 200 size-exclusion, and Mono Q ion-exchange chromatography. Purity (≥90%) was confirmed by Coomassie-stained SDS-PAGE (**Supplementary Fig. S3**) and LC-MS analysis of intact mass. Protein concentration was determined by Bradford assay using BSA as standard. The final preparation was buffer exchanged into 50 mM HEPES-NaOH pH 7.5, 150 mM NaCl, 1 mM TCEP, sterile-filtered (0.22 µm), and stored in aliquots at −80 °C. Freeze-thaw stability was verified, with no precipitation observed after two cycles. As an independent positive control, we also purchased a commercially available *E.coli*-derived SAE1/2 complex with no tags, which has 5 extra amino acids (GSPNS), on the N-terminus of SAE2 (R&D Systems, E-315).

### <u>E</u>lectrophoretic <u>M</u>obility <u>S</u>hift <u>A</u>ssay<u>s</u> (EMSAs)

Recombinant SAE1/2 from two sources (Genscript and R&D), or BSA (A7888-100G, Sigma) and Casein (Sigma-Aldrich 218682) as control was incubated with the various RNAs for 30 min at 25°C, in 50 mM HEPES-NaOH pH 7.5, 150 mM NaCl, 1 mM TCEP. Increasing molar ratios protein over RNA were used (20:1 to 200:1). Before electrophoresis, 1 μL of 50% v/v glycerol, was added to the reaction and complexes were resolved on 10% v/v native PAGE gels at 10 mAmps constant at 4°C for ∼3-4hrs. Gels were exposed overnight to Hyperfilm MP (Amersham 28-9068-45), at −80 °C without drying.

### Auto-SUMOylation of SAE2 (thioester formation)

5 pmol of recombinant SAE1/2 were mixed with 5-25 pmol of the SUMO1 peptide (R&D UL-712) and incubated for 4 min at 37 °C together with increasing concentrations of temp-refolded human 5’P-mascRNA (0-6 μM) (ThermoScientific R0441) and at 5 mM MgCl2. Reactions were initiated by adding ATP at 1 mM final concentration. A positive control (no mascRNA) and a negative control (no ATP) were also analyzed.

### *In vitro* SUMOylation assays

We utilized the *in vitro* SUMOylation (IVS) kit (Abcam ab139470) to perform IVS reactions and replaced the E1 enzyme of the kit with the custom made 6xHis-SAE1/2 from GenScript. Reactions were performed as per the manufacturer’s instructions with RANGAP1 fragment as protein substrate for SUMOylation (residues 418-587 of RanGAP1 fused to an N-terminal GST tag) and probed with the SUMO1 antibody of the kit. Negative control was a reaction with no addition of ATP. We confirmed the enzymatic activity of our SAE1/2 preparations by Western Blot detection of RANGAP1-SUMO1 product (see **Supplementary Fig. S3**).

### Cell culture, transfection, subcellular fractionation

293FT (ThermoFisher R70007) were cultured 37 °C under 5% CO2, in high glucose DMEM medium (Gibco 11965084) supplemented with 1% Glutamax (Gibco 35050061), Minimum Essential Amino-acids (MEM) (Gibco 1140050), Sodium Pyruvate (Gibco 11360070) and 10% v/v FBS (Atlas F0050D); hereafter, referred to as full growth medium. Media were filtered in 0.2 μm filter units (Fischerbrand FB12566502 or ThermoScientific 0974103). Trypsin/EDTA 0.05% (Gibco 25300054) was used to detach the cells from the culture vessel’s surface. Frozen stocks of cells were taken periodically using filtered freezing medium, composed of 90% full growth medium and 10% v/v DMSO. Cells from one T75 flask were trypsinized, collected, 1 mL of freezing medium added, and cells were stored in 2 mL cryovials in a liquid nitrogen tank.

### Knock-down of human SAE2 (UBA2)

Cells seeded in 6-well plates were transfected for 48 h with 25, 50 and 100 nM of UBA2 siRNA for silencing human UBA2 (SAE2) expression (Horizon Discovery J-005248-09-0005), using Lipofectamine-2000 (Invitrogen 11-668-019) for transfection following manufacturer’s instructions. For the 6 well plate, a first mixture was prepared by adding 10 μL Lipofectamine to 250 μL Optimem reduced serum, no phenol red (Gibco 11-058-021). A second mixture was prepared by adding 1 or 2 or 4 μL of 50 μM stock siRNA for UBA2 to 250 μL Optimem. The two mixtures were combined and incubated for 20 min at RT. The final mixture was added to full-growth medium 1.5 mL and the final volume 2 mL was added per well of the plate. Cells were collected for protein analysis at 48 h post transfection, supplemented with protease/phosphatase inhibitors (Halt™ Protease and Phosphatase Inhibitor Cocktail 100X - ThermoFisher PI78442), sonicated and centrifuged at 4°C for 15 min (on a Sorvall ST 8 centrifuge). The supernatant was mixed with 1:1 v/v reducing loading buffer (4xLDS NuPAGE buffer (Invitrogen NP0007) + 1/10 volume DTT 0.5 M), then heated at 70°C for 12 min, before analyzing on SDS PAGE.

Synthetic human mascRNA and synthetic scrambled control RNA oligonucleotides (synthesized by IDT, Reverse-Phase HPLC-purified, sequences shown below) were temperature-refolded (95 °C to 25 °C) and were transfected in a forward manner, at 0.1-10 nM final concentration using Lipofectamine 2000, for up to 48hr in plated 293FT. For a 10 cm cell culture dish, two parts were first prepared: a) 50 μL Lipofectamine was added in 1250 μL Optimem and b) 1 μL of refolded 100 μM stock RNA oligonucleotide with 1250 μL Optimem. Next, these two parts were mixed, and complexes were allowed to form for 20 min at RT, inside the biosafety cabinet and added to 7.5 mL of growth medium without antibiotics (total = 10mL medium). For 8-well culture chambers, transfections were scaled down proportionally to 250 μL final volume per well.

RNA oligonucleotides for transfection experiments:

#### mascRNA

5’-P-GAUGCUGGUGGUUGGCACUCCUGGUUUCCAGGACGGGGUUCAAAUCCCUGCGGCGUCU

#### scrambled-mascRNA

5’-P-GCGUGCCCCGAUGCAGGCUGUAGCGGCGUGGUUAACAGGGUCCUUUCGUUUCGCUGUA

The plasmid pCDNA5/FRT/TO 6XHis-FLAG SUMO1/2/3 WT, was transfected using the Turbofect reagent (ThermoScientific FERR0531) at 10-25 μg quantities per 10 cm dish, in the same manner as Lipofectamine.

### Biochemical cell fractionation

For cellular fractionation, the REAP protocol was used (36). Briefly, cells were collected in 1000 μL 1x cold PBS and separated into 2 parts. Part A, 100 μL of the cell suspension, was used to prepare the whole cell lysate (WCL); the cells were pelleted by a brief centrifugation before lysis buffer addition. Part B, 900 μL of the cell suspension, was pelleted and resuspended in 0.1% v/v NP-40 in 1x PBS, and centrifuged at 1000 x g for 1 min at 4°C. The supernatant was collected as the cytoplasmic fraction and the pellet was washed one time with 0.1% v/v NP-40 in 1x PBS and collected with 1 min centrifugation at 1000 x g. The cell pellet for WCL and the nuclear pellet were lysed in RSB150: 150 mM NaCl, 50 mM Tris pH 7.5, 0.5% IGEPAL, 0.1% Triton X-100, 1.25 mM EDTA, supplemented with protease, phosphatase (Thermo 354118) and peptidases inhibitors, sonicated as described above. Centrifuged and cleared lysates were mixed with 1:1 v/v sample buffer, containing NuPAGE 4xLDS (Invitrogen NP0007) +1/10 DTT 0.5 M (Roche 10278200). Finally, all samples were placed at 70 °C for 12 min. We also used a commercial cell fractionation kit, following the manufacturer’s instructions (Thermofisher 78840).

### Immunoprecipitations (IPs), Western blotting

IP was performed using Dynabeads A (Invitrogen 10002D). We bound 6 μg of mouse polyclonal Ab (anti-SAE2: Bethyl A302-925A; anti-SUMO1 Abcam (ab32058) or 6 μg of Normal Rabbit Serum (NRS) or rabbit IgG (Genscript A01008) on beads resuspended in 1x PBS, 0.1% v/v Nonidet P40 (IGEPAL) (Alfa Aesar J61055) for 1 h at room-temperature. Briefly, cells were trypsinized, detached and washed with PBS, which was discarded before lysis. We used 9μL volume of lysis buffer supplemented with protease and phosphatase inhibitors (ThermoScientific 78442) per mg of pelleted cells, and cells were lysed by sonication. For inhibiting de-SUMOylation and enhancing its detection, the isopeptidases inhibitor NEM (N-ethyl maleimide) was added in the lysis buffer to final 20 mM. For isolating RNA from the beads-bound protein, 10 μL of rRNasin (Promega N2511) was added per 400 μL of cell lysis as described above to prevent RNA degradation. Lysis buffer was slowly added onto the frozen cell pellet and pipetted carefully to allow for complete dissociation of the cells. Cell lysate was sonicated with 15 pulses on sonicator (Sonifier 450, Branson) with 40% duty cycle. Following this, cells were centrifuged at 17000 x g, for 15 min, at 4 °C and the supernatant was incubated with the Ab-beads for 5 min for SAE2 IP, 3 hr for SUMO1 IP. Proteins were eluted from the IP beads 30 μL of 1/3 RSB-150+ 2/3 4xLDS+ 1/10 β-mercaptoethanol (or DTT) sample buffer and placed at 70 °C for 12 min. Samples were run on 4-12% m/v Bis-Tris NuPAGE gels (Invitrogen, NP0322BOX) and transferred on nitrocellulose (Cytiva, 10600001). Protein ladders iBright (Thermo Fischer LC5615) and PageRuler (Thermo Scientific 26619) were used to evaluate MWs. Single or dual fluorescence detection of protein bands was performed on a ChemiDoc MP system (Bio-Rad). Densitometric analysis of immunoreactive bands was determined using Bio-Rad Image Lab software, or ImageJ software.

### RNA-protein pull-down assay for proteomics

To identify mascRNA-interacting proteins, we bound the biotinylated RNA oligos on MyOne Streptavidin C1 beads (Invitrogen 65001), following the manufacturer’s protocol. RNA oligonucleotide baits with 3’ biotin-TEG purified by RP-HPLC (IDT) were used. For one pull-down, 50 μL of streptavidin beads slurry were washed consecutively with 1 mL 1x Solution A (NaOH 0.1M, NaCl 0.05 M), 1 mL of 0.1 M NaCl, and 3x with 1 mL B+W solution (10 mM Tris HCl pH 7.5, 1 mM EDTA, 2 M NaCl, 0.05% v/v Tween-20) and resuspended in 100 μL of the same buffer 2 μL (100 pmol / μL) of temperature-refolded biotinylated RNA oligo (200 pmol total) were added and incubated for 15 min at RT under rotation. Following this, beads were washed 3x with RSB-150 and then mixed with a cell lysate prepared as for IP, and incubated for 1 h at 4 °C, under rotation. Beads were washed 3x with 1 mL of RSB-150. For Western blot, proteins were eluted by adding 30 μL of 1/3 v/v RSB-150 + 2/3 v/v 4x LDS + 1/10 β-mercaptoethanol sample buffer and placed at 70 °C for 12 min.

Biotinylated RNA probes:

#### scrambled-mascRNA

GCGUGCCCCGAUGCAGGCUGUAGCGGCGUGGUUAACAGGGUCCUUUCGUUUCGCUGUA-3’ biotinTEG

#### mascRNA

5’P-GAUGCUGGUGGUUGGCACUCCUGGUUUCCAGGACGGGGUUCAAAUCCCUGCGGCGUCU-3’ biotinTEG

#### mascRNA-CCA

5’P-GAUGCUGGUGGUUGGCACUCCUGGUUUCCAGGACGGGGUUCAAAUCCCUGCGGCGUCUCCA-3’ biotinTEG

### Sample preparation for proteomic analyses

For identifying mascRNA and SAE2-associating proteins we performed SAE2 IP and mascRNA pull down respectively, and SUMO1 IP for identifying sumoylated proteins, as described above. From these experiments, proteins were eluted from Dynabeads A, using 30 μl of 6 M Urea, 100 mM Tris-HCl pH 7.5, for 10 min, at 30 °C and 1000 rpm. A fraction of 5 μL from the 30 μL eluate were used for WB and 7 μL for SYPRO or Coommassie stain. The remaining ∼18 μL of sample were analyzed by mass spectrometry at Taplin Mass Spectrometry facility, Harvard University.

### RNA isolation

Total RNA from cells was isolated with miRvana kit (Ambion AM1560) following manufacturer’s instructions. RNA from IP beads was isolated using Trizol (Invitrogen 10296010) following manufacturer’s instructions. Before Northern or RT-qPCR, RNA was treated with RQ1 DNase (Promega M6101) followed by Phenol/Chloroform extraction EtOH precipitation. Briefly, DNase reactions were brought to 400 μL of volume by adding water, and equal volume of Phenol/Chloroform/Isoamyl alcohol 25:24:1 (Invitrogen 15593-031) was added. After vortexing (30 sec) and centrifugation (5 min, 12000 x g, RT) the top (aqueous) phase was retrieved. The process was repeated with Chloroform/Isoamyl 24:1 (Millipore-Sigma 25666) to remove traces of phenol. RNA was precipitated from the water phase by adding: 2.5x volume of ice cold EtOH, 1/10x volume 3M Sodium Acetate pH 5.5 (Invitrogen AM9740), and 5 μg of glycogen (Invitrogen AM9510), and incubating for 30 min at −80 °C followed by centrifugation at 17, 000 x g 4 °C for 30 min, washed with ice cold 70% EtOH, centrifuged as before, and air dried for 5 min. The RNA pellet was resuspended in 15 μL nuclease-free water.

### cDNA synthesis (reverse transcription)

The miScript kit (Qiagen 218161) was used for reverse transcription, following manufacturer’s instructions. For RNA extractions following immunoprecipitations only, we added 11 μL input of RNA and 1 μL (corresponding to 1 ng) of *Renilla luciferase* mRNA in vitro transcription product (using the Renilla Luciferase plasmid, rLuc Promega E2231) as spike-in RNA for normalization. For whole-cell RNA, the miScript protocol was followed without the addition of spike-in RNA.

### qPCR

qPCR was performed in 25 μL final volume using the cDNAs generated above. 12.5 μL of 2x SYBR Green PCR Master Mix (Applied Biosystems 43-091-55) was mixed with 2 μL cDNA template (5 ng/μL), 300 nM (final) of each forward and reverse primer and nuclease-free water to a final volume of 25 μL. Reactions were run on QuantStudio 3 Real-Time PCR System (ThermoScientific A28567) for 40 cycles. Results were analyzed with an integrated software QuantStudio Design and Analysis (Applied Biosystems), using the Comparative Cт (ΔΔCт) method. For the RNA IPs, rabbit or mouse IgG-retrieved RNA served as control sample and spike-in luciferase RNA as reference gene for normalization of target RNA levels. For whole cell RNAs, GAPDH was used for normalization of target gene expression.

*MALAT1* Fwd:

GCTGACACTTCTCTTGACCTTAG; *MALAT1* Rev: CACCGAAGGCTTAAAGTAGGAC (Hs.PT.58.26451167.g IDT); mascRNA Fwd: GATGCTGGTGGTTGGCACTC; mascRNA Rev: TGGAGACGCCGCAGGGAT (from Sun *et* al, 2021); Pair no2 mascRNA fwd: CTGGTGGTTGGCACTCCT; mascRNA rev: AGACGCCGCAGGGATTTG; *GAPDH* Fwd: ACCCACTCCTCCACCTTTGAC; *GAPH* Rev: ACCCTGTTGCTGTAGCCAAATT; Luciferase Fwd: CGCTGAAAGTGTAGTAGATGTG; Luciferase Rev: TCCACGAAGAAGTTATTCTCCA.

### Northern Blot

RNA samples mixed 1:1 v/v in formamide-containing RNA loading dye, heated at 70°C for 1 min and cooled to 4°C. 1 μg of total RNA was analyzed on 10-15% Urea-PAGE in 1x TBE at 10 mA. The RNA species were transferred onto a Nylon membrane (positively charged) (Roche) in 1x TBE for 15 min at 20 V constant, in a semi-dry apparatus, using 1x TBE presoaked blotting papers (3) below and (3) on top of membrane-gel assembly. The blot was soaked for 2 min in 1x TBE buffer. RNA was then UV-crosslinked (immobilized) to the membrane (1200 μJoules). Membrane was preincubated in 5 mL Church buffer (for 50 mL: 36 mL 1 M Na2HPO4, 14 mL 1 M NaH2PO4, 0.2 mL 0.5 M EDTA and 3 mL 20% v/v SDS), then hybridized in fresh Church buffer with 0.2 nM 5’ DIG(Digoxigenin)-probes antisense to either mascRNA or U6snRNA in a hybridization chamber with rotation for 1 h at 55 °C (sequences of RNase-free HPLC purified DNA Northern probes: Human mascRNA /5DigN/AG AC+G CCG CAG GGA +TTT GAA CCC CGT CCT GGA AAC CAG GAG TGC CA+A CCA CCA GC+A TC (58 nt), Human U6snRNA /5DigN/CA C+GA A+TT TG+C GT+G TCA TC+CTT (22 nt), where + denotes <u>L</u>ocked <u>N</u>ucleic <u>A</u>cid (LNA) modification.

Following hybridization, 10 min washes were performed sequentially with 3 mL of 2x Saline-Sodium Citrate Buffer (SSC) / 0.1% v/v SDS, 1x SSC / 0.1% v/v SDS, 0.2x SSC / 0.1% v/v SDS (by diluting 20x SSC, Invitrogen AM9770) at hybridization temperature. The blot was then placed in 20 mL of 1x digoxigenin (DIG) wash buffer for 15 min at room temperature, followed by 30 min in 1x DIG blocking buffer. Anti-digoxigenin-Alkaline Phosphatase (AP) antibody was added in 1x DIG blocking buffer for 30 min at room temperature. Blot was washed twice in 1x DIG Wash buffer for 15 min and then equilibrated in 1x DIG detection buffer. Visualization was performed with a chemiluminescent substrate of AP (CDP-Star; Merck C0712). U6snRNA probing was performed on the same blot, after washing previous CDP-star.

### Immunofluorescent staining and confocal microscopy

293FT cells were seeded onto 8-well chamber slides (Falcon 354118) precoated with poly-D-lysine (Gibco A3890401). 250 μL of cell suspension in full-growth medium were seeded in each well and let to adhere for 6 h. Cells were then washed with 1x PBS and fixed with 200 μL of 4% v/v Paraformaldehyde (PFA) (Biotium 22023) for 20 min, at room temperature. PFA was removed and cells were washed 3 times with 1x PBS. Cells were permeabilized with 200 μL of 0.1% v/v Triton and 1% m/v BSA in 1x PBS for 20 min, at room temperature. Following permeabilization, cells were washed three times with 200 μL 1x PBS. Blocking was performed with 200 μL 3% m/v BSA in 1x PBS for 20 min at room temperature. After blocking, three washes in 1x PBS were performed. Chambers were stained with SAE2 (Bethyl A302-925A), or SUMO1 (Santa Cruz sc-5308) primary antibodies 1/1000, overnight at +4 °C, on slow motion (∼10 rpm) inside a humidified box, protected from light. The next day, the primary antibody solution was removed, and wells were washed 3 times with 1x PBS for 5 min each. A 200 μL mixture of secondary antibodies (anti-rabbit (Invitrogen A21245), anti-mouse (Invitrogen PIA32766TR) and DAPI (Thermo PI62247) at 0.5 ng/mL was added per well and incubated for 1 h at room temperature. Each well was then washed 3 times, with 200 μL 1x PBS, the chamber walls were removed and the slide was air dried. A drop of anti-fade Mounting medium (Prolong Antifade mounting medium (Invitrogen P36931) was added on each well and a coverslip (Fischerbrand 12-545J) was placed on top of wells. For stacks and image acquisitions, the microscope Olympus IX83 P2ZF was used with objective length UPLAPO OHR 100x / 1.50 numerical aperture and Multicamera (Hamamatsu ORCA-fusion) and operated with Software OLYMPUS cellSens Dimension 3.2, 23706. The channels used were: DAPI, GFP, mCherry and Cy5.

### Bioimage analysis

Surfaces (μm^2^) and numbers of objects were quantified, on channels thresholded at the same intensity value between experimental conditions (i.e. control oligo vs mascRNA transfection), using the 3D objects counter tool in the ImageJ application. Moreover, colocalization was quantified using either Imaris colocalization or ImageJ colocalization tools JACoP. At least 6 fields of view (FOVs) per experimental condition per image were used for analysis.

### Data analysis

Densitometric analyses were performed using ImageLab (Biorad), version 6.1.0, build 7, standard edition. Standard t-test was used for pair-wise comparisons, statistical significance was set at p≤ 0.05, and data were presented with GraphPad Prism, v8.

### Software tools

NLS mapper (37) (https://nls-mapper.iab.keio.ac.jp/cgi-bin/NLS_Mapper_form.cgi), NoLS mapper (38) (https://www.compbio.dundee.ac.uk/www-nod/index.jsp), IUPRED3 (39) (https://iupred3.elte.hu/) used to assess localization signals and disordered regions in investigated proteins. PyMOL (educational license) was used to depict mascRNA model and protein regions.

## RESULTS

### mascRNA adopts a 2D cloverleaf and 3D L-shape fold in solution

The conformation of the mascRNA transcript in solution was initially investigated using enzymatic and chemical probing. RNases T1 and S1, together with lead acetate, were used to assess the accessibility of single-stranded regions, while RNase V1 was employed to detect double-stranded or structured regions. **Fig. 1A** summarizes the probing results, revealing moderate cleavage by enzymatic probes but extensive cleavage with lead acetate. The analysis clearly highlighted the three loops of a cloverleaf structure. Strong cleavages by RNases S1 and T1 were observed at positions 14 and 26-28, along with moderate cleavages in neighboring regions, including nucleotides 40-42. These same sites were also cleaved by lead acetate, suggesting the presence of a 7-nucleotide loop in region II (D arm equivalent), a 5-nucleotide loop in region III (anticodon arm equivalent) and a 7-nucleotide loop in region IV (T arm equivalent). Region I (acceptor stem equivalent) was not accessible to this analysis, but is very likely to be double stranded as seen in the crystal structure (3) and in our SAXS model (see below). Regions II and III show a mixed reactivity for probes specific for single and double stranded RNA. For instance, both lead and RNase V1 cleavages are observed for residues 19-22, suggesting some flexibility and the possibility of alternate conformations. This dynamical behavior is also illustrated by the difference between the 2D structure deduced from the probing and that seen in the crystal structure, as well as in the model described below, with a loop of 7 vs 5 nucleotides and a stem of 2 vs 3 base pairs, respectively. In contrast, region IV contains four consecutive guanosines (35–38), which were resistant to RNase T1 under both native and denaturing conditions (see G-lane with T1 at 60 °C), indicating their involvement into a very stable double-stranded structure of 5 base pairs. This is corroborated by the absence of lead acetate cleavage in the same region. Overall, probing data strongly support that mascRNA in solution adopts a cloverleaf structure (**Fig. 1B**).

**Figure 1.**
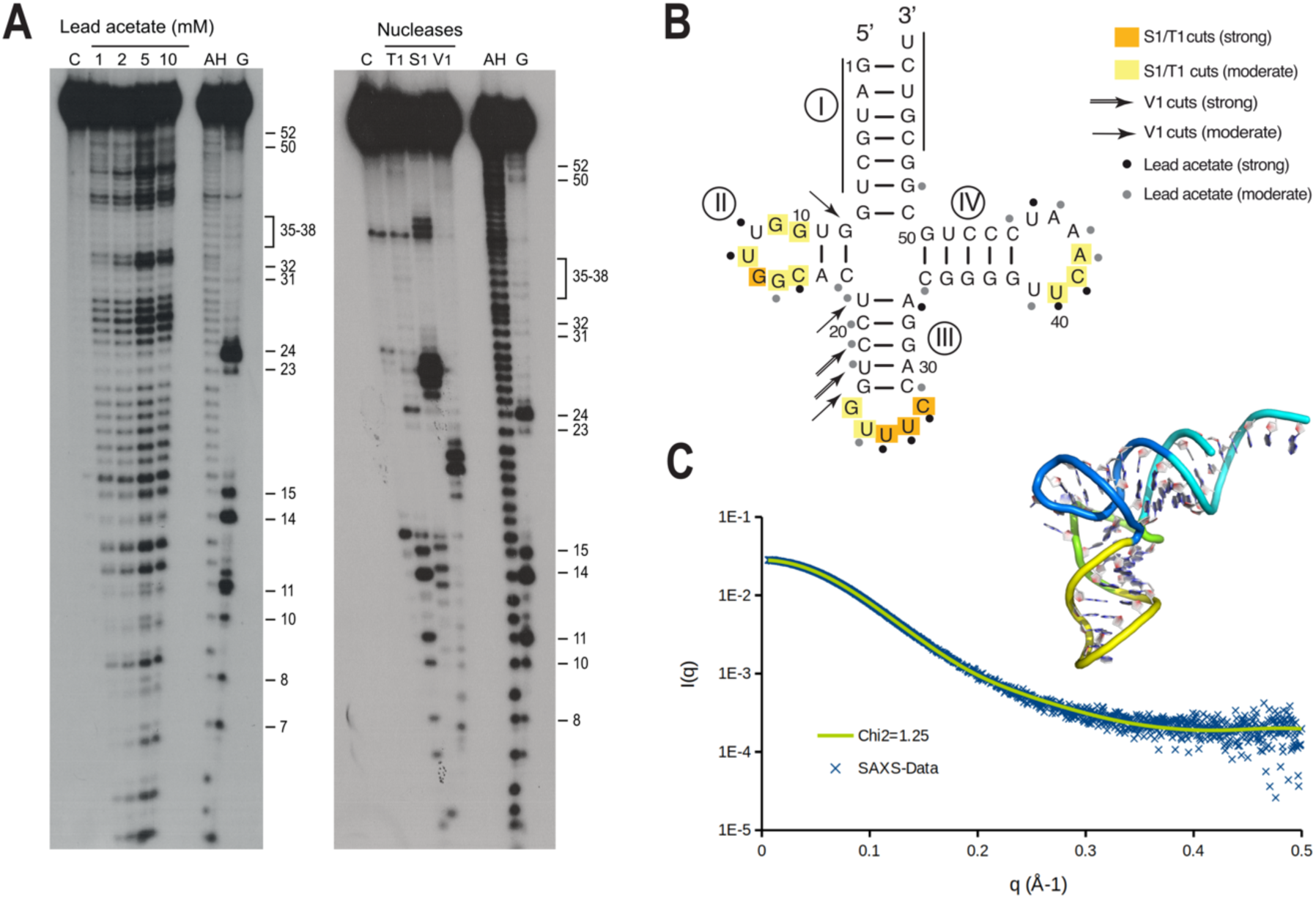
Human mascRNA adopts tRNA-like fold. **(A)** Autoradiograms of a 14% m/v denaturing polyacrylamide gel showing probing experiments performed on radiolabeled mascRNA using lead acetate (left panel) and nucleases T1, S1, and V1 (right panel). Control reactions (lane C) were conducted in parallel without probes. AH: alkaline hydrolysis; G: denaturing RNase T1 digestion, with G residues indicated. **(B)** Summary of the probing results mapped onto the mascRNA 2D structure (as seen in the model below). Four regions of interest (I, II, III, and IV) are highlighted. Bars at the 5’ and 3’ ends mark regions that could not be resolved. **(C)** SAXS analysis and modeling of mascRNA showing a good agreement (Chi2 = 1.25) of the theoretical curve (in green) calculated from the model with the experimental data (in blue). The four regions (I, II, III, and IV) of mascRNA are depicted in cyan, green, yellow and blue, respectively.

The analysis of mascRNA transcript by SEC-SAXS revealed a radius of gyration *Rg* of 20.2 Å and a maximum distance *Dmax* of 72 Å (**Supplementary Fig. S1A-D**), which are smaller than those reported for conventional tRNAs (Rg = 24 Å, Dmax = 81 Å) (40, 41). Several atomic models built with different software and web services were evaluated. The best fit to the experimental SAXS data (**Fig. 1C**) was obtained with a model generated by trRosettaRNA using a multiple sequence alignment of 6 mascRNA sequences (from human, mouse, chimp, rhesus monkey, lizard and *Xenopus tropicalis*). This model achieved a goodness-of-fit (Chi^2^) of 1.25, substantially better than the crystal structure (PDB ID 8v1i (3) chain A / B: Chi^2^ = 9.2 / 10.4) or the AlphaFold3 model (Chi^2^ = 4.5). Nevertheless, the trRosettaRNA model exhibits highly similar secondary and tertiary folds, with root-mean-square deviations (RMSD) of 3.1 Å and 3.4 Å relative to chains A and B of the crystal structure (intra-chain RMSD = 1.9 Å for A/B alignment over 56 nucleotides). Taken together, our data support the folding of mascRNA into a cloverleaf-like secondary structure and a 3D L-shape fold in solution.

### mascRNA interacts with the SUMO E1 enzyme in vitro and in cellulo

To determine whether mascRNA associates with the SUMO E1 enzyme in cells, we immunoprecipitated endogenous SAE2 using a custom-generated anti-SAE2 antibody (see Ab validation in **Supplementary Fig. S2A and B**) and quantified co-purifying mascRNA by RT-qPCR. This analysis revealed a significant enrichment of mascRNA in SAE2 immunoprecipitates compared to control IgG IPs, establishing an in-cell association between mascRNA and SAE2 (**Fig. 2A and B**). We next explored whether mascRNA directly contacts the SUMO E1 complex *in vitro* and where on the RNA such binding occurs. For these experiments we acquired highly pure and catalytically active N-terminally 6xHis-tagged recombinant SAE1/2 complex (**Supplementary Fig. S3A-C**). We performed electrophoretic mobility shift assays (EMSAs) with 5’ radiolabeled mascRNA and increasing concentrations of recombinant SAE1/2 (**Fig. 2C**), which revealed a clear, dose-dependent mobility shift, demonstrating the formation of a stable RNA-protein complex. We confirmed this result with purified recombinant SAE1/2 without 6xHis from a different source (**Supplementary Fig. S3D**), and we also observed only partial competition of mascRNA-SAE1/2 complex formation by excess total yeast RNA (**Fig. 2D**, **Supplementary Fig. S3E**). In addition, RNase S1 foot-printing of mascRNA in the presence of SAE1/2 versus BSA revealed a protection pattern specifically at the anticodon loop equivalent, consistent with a direct RNA-protein contact (**Fig. 2E**). These experiments revealed a selective and specific association between mascRNA and the SUMO E1 complex.

**Figure 2.**
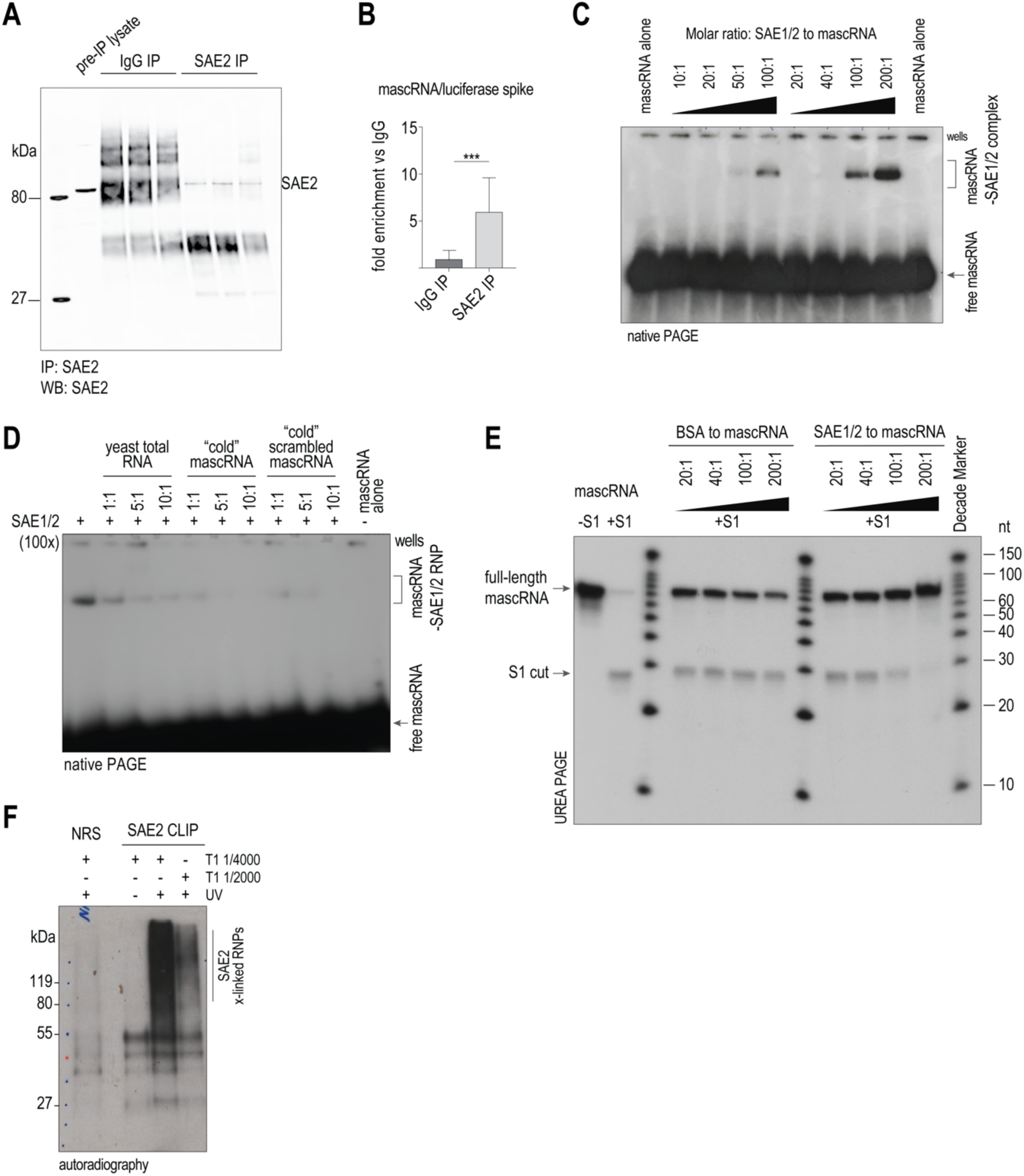
Human mascRNA associates with SUMO E1 heterodimer enzyme *in vitro* and in cellulo. (**A**) SAE2 IP and immunoblot, IgG was used as a negative control. (**B**) RT-qPCR analysis of RNAs co-purified with endogenous SAE2 immunoprecipitates compared to IgG control IPs, showing significant enrichment of mascRNA. Luciferase spike-in RNA was used for normalization. Data are shown as means ± S.D, *n* = 3 vs 3 independent IPs, ***p = 0.007. (**C**) Electrophoretic mobility shift assays (EMSAs) of 5’ radiolabeled mascRNA incubated with increasing concentrations of purified SAE1/2 heterodimer. The SAE1/2 complex with mascRNA is indicated. (**D**) EMSA of mascRNA and SAE1/2 in the presence of increasing amounts of indicated unlabeled RNA species, to test the SAE1/2-mascRNA complex specificity. (**E**) RNase S1 footprinting of mascRNA in the presence of purified recombinant SAE1/SAE2 heterodimer compared to BSA. A predominant protection pattern at the anticodon loop equivalent indicates direct binding of mascRNA to the SUMO E1 complex. Electrophoretic mobility shift assays (EMSAs) of 5’ radiolabeled mascRNA incubated with increasing concentrations of purified SAE1/2 heterodimer. The SAE1/2 complex with mascRNA is indicated.(**F**) Crosslinking and immunoprecipitation (CLIP) with anti-SAE2 antibody under high-stringency lysis conditions (2% Empigen), showing robust RNA crosslinking to SAE2. NRS IP, and no-UV crosslinking of cells were used as negative controls.

Because these *in vitro* assays support a direct interaction, we next asked whether SAE1/2 engages RNA more broadly *in vivo*. The anti-SAE1 antibodies that we tested did not immunoprecipitate SAE1 efficiently, so we focused on SAE2. UV crosslinking and immunoprecipitation (CLIP) with our anti-SAE2 antibody under stringent lysis conditions (2% Empigen) revealed robust radioactive RNA signals in SAE2 immunoprecipitates but not in IgG or non-crosslinked controls (**Fig. 2F**). Notably, only SAE2 and not SAE1 was recovered in the CLIP immunoblots (**Supplementary Fig. S3F**), indicating that the stringent lysis conditions disrupted protein-protein associations, thus validating a direct interaction of RNA with SAE2. Thus, these results demonstrate that, at a minimum, SAE2 directly contacts RNA *in vivo*, while not precluding RNA association by SAE1.

Finally, to assess whether mascRNA alters SUMO E1 integrity or function, we tested SAE1/2 complex formation and enzymatic activity *in vitro* in the presence of mascRNA. MascRNA did not affect complex stability in native PAGE analyses, nor did it alter E1 activity in assays measuring SAE2 auto-SUMOylation or RANGAP1-SUMO1 conjugation (**Supplementary Fig. S4A-C**). These results indicate that mascRNA engages the SUMO E1 complex without disrupting its enzymatic activity.

Together, these complementary approaches demonstrate that mascRNA directly associates with the SUMO E1 enzyme *in vitro* and in cells, establishing a specific RNA-protein interaction that links mascRNA to the SUMOylation pathway.

### Nuclear localization of SAE2 is modulated by exogenously provided mascRNA

We next sought to define the cellular context of SAE2 localization and function. As a first step, we performed proteomic analysis of SAE2 immunoprecipitates (**Fig. 3A and B**), aiming to map its interaction network and uncover mechanisms that could be influenced by mascRNA. Mass spectrometry confirmed specific retrieval of SAE2 and its known partner SAE1 (**Fig. 3B**), and identified a set of 43 SAE2-associated proteins (0 peptides in normal serum IP, ≥2 peptides in SAE2 IP) enriched for nuclear localization (many linked to nuclear speckles and nucleolar processes), RNA-binding functions, histone-modifying enzymes and a smaller subset of plasma membrane and cell junction proteins (**Fig. 3B**, **Supplementary Fig. S5, Supplementary Table S1**). STRING network analysis supported the statistical significance of these findings, showing that the majority of SAE2-enriched proteins were annotated as nuclear (37/44, FDR: 4.06×10^-8^), many of which are involved in histone acetylation (8/44, FDR: 4.19×10^-12^), or are RNA binding factors (21/44, FDR: 2.88×10^-8^) involved in post-transcriptional regulation (**Supplementary Fig. S5**). Together, these proteomic data position SAE2 at the intersection of nuclear regulatory pathways.

**Figure 3.**
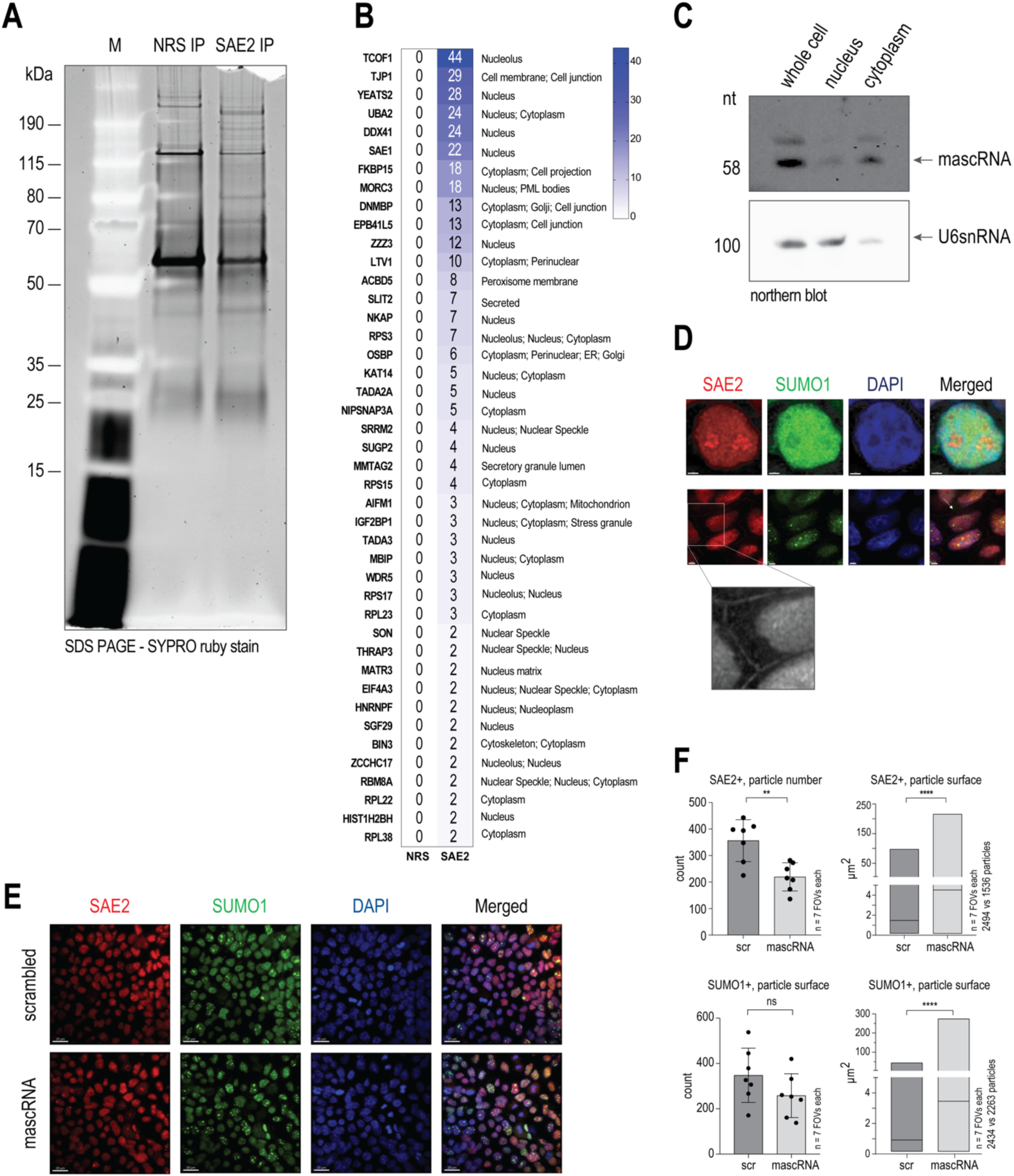
SAE2 nuclear localization and its modulation by mascRNA **(A).** SYPRO Ruby stain of urea-eluted proteins from NRS and SAE2-IPs. (**B**). SAE2-IP proteomics vs NRS control in untreated cells. Heatmap of unique peptide number per identified protein per IP and subcellular localization (Uniprot). (**C**). Cellular fractionation and mascRNA northern blot from nuclear and cytoplasmic fractions; U6 was probed as a control of the fractionation. (**D**). SAE2 and SUMO1 immunofluorescence of untreated cells. Scale bar: top row 2 μm, bottom 3 μm. (**E**). mascRNA and scrambled oligo treatment of 293FT cells, SAE2 and SUMO1 immunofluorescence. Scale bar: 20 μm. (**F**). Quantifications and statistic test of surface area of SAE2^+^ signal and SAE2^+^ particle numbers after scrambled oligo control (scr) and mascRNA treatments. Data are shown as means ± S.D; Student’s *t*-test \*\**p*=0.0025, \*\*\*\**p*<0.0001, SUMO1: n.s, *p*=0.1493, *n*=7 FOV per condition.

To determine whether mascRNA itself is present in the same cellular compartments as SAE2, we performed biochemical cell fractionation followed by Northern blot analysis. The majority of endogenous mascRNA was detected in the cytoplasm; however, a distinct fraction was present in the nucleus (**Fig. 3C**), quite possibly representing the mature mascRNA species processed from *MALAT1*.

We next examined SAE2 localization in cells by immunofluorescence (IF). In control cells, SAE2 showed distinct nuclear distributions, in agreement with the presence of predicted C-terminal nuclear localization signals in SAE2 amino acid sequence (19). Within the nucleus, SAE2 displayed both diffuse and punctate signals, with condensate-like structures detected in DAPI-depleted regions, a hallmark of nucleolar localization (**Fig. 3D**). A weaker signal was also present at the cell periphery, localized in potential membrane-associated pools (**Fig. 3D**, bottom row). SUMO1 immunostaining displayed a comparable diffuse and punctate pattern that was predominantly nuclear, though spatially distinct from SAE2, with only partial colocalization (**Fig. 3E**).

Finally, we asked whether mascRNA affects SAE2 and SUMO1 distributions *in situ*. Upon transfection with synthetic mascRNA, the SAE2 punctate signal was markedly altered: the overall SAE2-positive area per field of view increased, while the number of discrete SAE2-positive particles decreased compared to a scrambled oligo control of identical length and nucleotide composition (**Fig. 3E and F**). Parallel analysis of SUMO1 revealed a significant expansion of the nuclear punctate surface, accompanied by a trend toward fewer SUMO1-positive foci in mascRNA treated cells (**Fig. 3E and F**).

These results indicate that mascRNA modulates both SAE2 and SUMO1 organization in the nucleus, thereby linking the observed mascRNA-SAE1/2 interaction with downstream changes in SUMO-related nuclear architecture.

### mascRNA associates with nuclear proteins, many of which are SUMOylated

We next investigated the broader protein network associated with mascRNA. Biotinylated RNA pull-down assays were performed using three synthetic baits: a scrambled mascRNA control, a mascRNA and a mascRNA-CCA oligo. Mass spectrometry revealed comparable unique and total peptide numbers across the three conditions (5159 for scrambled oligo, 5594 for mascRNA, and 5331 for mascRNA-CCA), enabling direct comparison of enrichment patterns (**Fig. 4A and B**). Among the 164 proteins identified with ≥2 peptides in any one of the two mascRNA pull-downs and absent (0 peptides) from the scrambled control (**Supplementary Table S2**), 143 carried a nuclear annotation (FDR = 1.27×10^-34^) and 39 were specifically annotated as nucleolar (FDR = 8.80×10^-14^) (**Fig. 4C**). RNA and rRNA metabolic process (FDR: 5.67×10^-34^ and 4.07×10^-9^ respectively) were also highly enriched. STRING network analysis revealed that these proteins formed a densely interconnected cluster (PPI enrichment p < 1.0×10^-16^), and notably, 111 out of 164 are known SUMOylation targets (42) (**Fig. 4C**).

**Figure 4.**
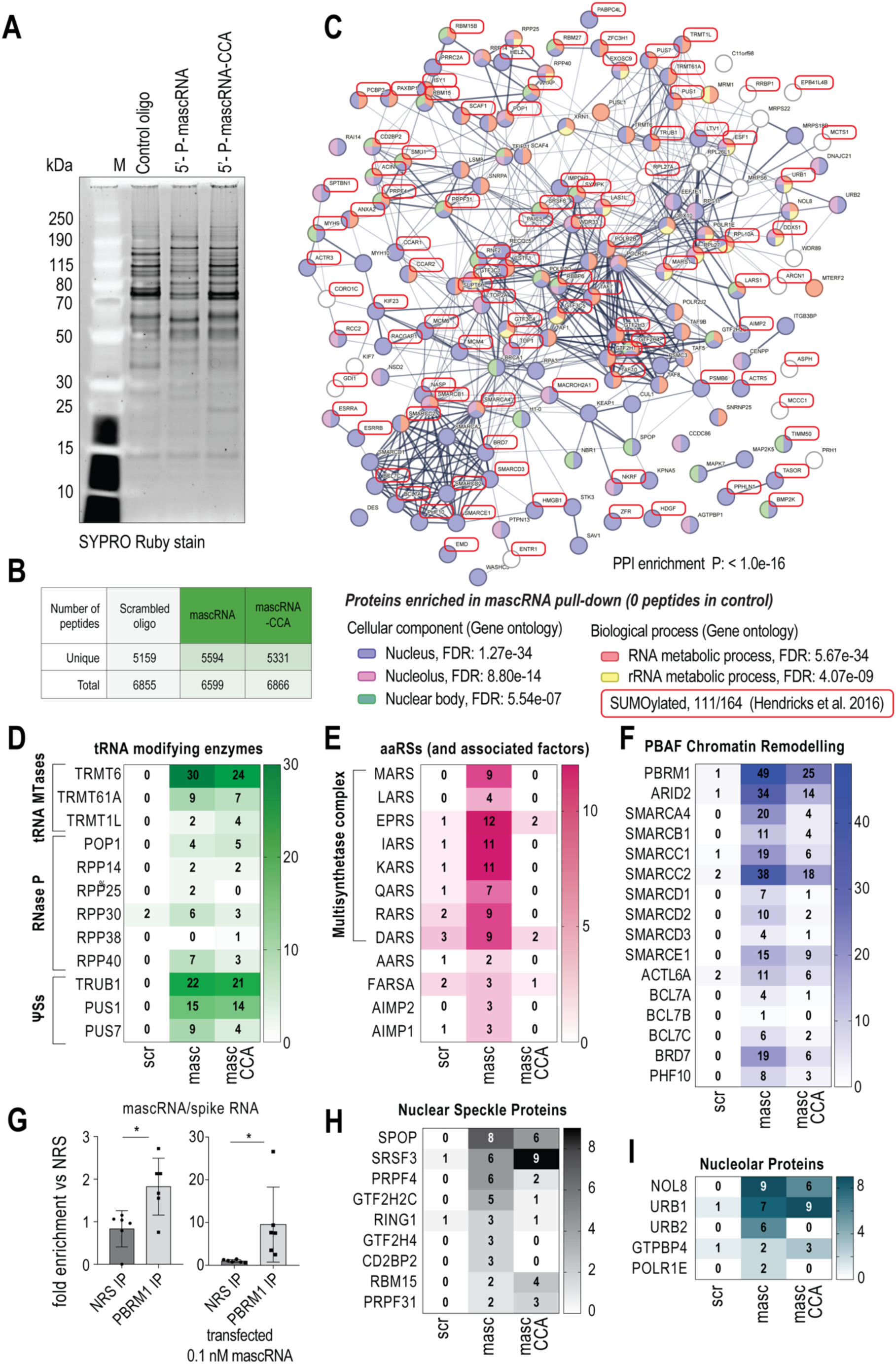
MascRNA interacts extensively with nuclear factors. (**A**). SDS-PAGE analysis of proteins eluted from biotin-RNA pull down experiments with the indicated oligos. (**B**). Number of peptides identified in biotin-RNA pull down experiments. (**C**). STRING analysis for proteins enriched in mascRNA pull-down (≥2 peptides in either mascRNA or mascRNA-CCA pull down and 0 peptides in scrambled control). Select enriched terms are shown, and proteins are colored for the annotated terms as indicated. Proteins experimentally identified as SUMOylated are highlighted in red. (**D**). tRNA modifying enzymes identified in biotin-RNA pull downs. In this and panels E, F, H and I, slightly different criteria were applied to uncover enriched retrieval in mascRNA pull-downs of various subunits of complexes or factors in the same category (≤2 peptides in control and more peptides in either mascRNA vs control pull down). (**E**). and (**F**). aaRSs and PBAF chromatin remodeling factors identified in the biotin-RNA pull downs. (**G**). Detection of endogenous and transfected mascRNA by RT-qPCR (6 technical replicates) in PBRM1 RNA-IP (2 replicate IPs). Data are shown as mean ± S.D, Student’s *t*-test. endogenous mascRNA \**p*=0.0120, endogenous and synthetic \**p*=0.0384. Luciferase levels: n.s. Data are shown as *n*=2 IPs, per condition. (**H**) and (**I**). Nuclear speckle (H) and nucleolar proteins (I) identified in biotin RNA pull downs.

Both mascRNA and mascRNA-CCA pull downs showed robust enrichment of factors involved in tRNA processing and modification, including subunits of the RNase P/MRP complex, pseudouridine synthases (TRUB1, PUS1, PUS7), and methyltransferases such as TRMT6/TRMT61A and TRMT1L (**Fig. 4D**). These associations validate the tRNA-like structural configuration of mascRNA, as it is recognized by canonical tRNA-interacting enzymes.

In addition, multiple aminoacyl-tRNA synthetases (aaRSs) belonging to the MultiSynthetase Complex (MSC) were detected, particularly with the mascRNA bait (**Fig. 4E**), in agreement with earlier reports (3, 8). Few aaRS peptides were also detected in the scrambled oligo control. Enrichment was substantially reduced in the mascRNA-CCA pull-down, suggesting that the presence of a 3′CCA end may modulate these interactions.

Beyond tRNA-modifying enzymes, mascRNA was associated with nuclear and chromatin-remodeling factors. Subunits of the Polybromo-associated BRG1-associated factor PBAF complex (PBRM1, ARID2, SMARCA4, among others) were specifically retrieved, with enrichment stronger in the mascRNA pull-down compared to mascRNA-CCA (**Fig. 4F**). This interaction was independently validated by PBRM1 immunoprecipitation, which recovered both endogenous and transfected mascRNA as measured by RT-qPCR relative to a luciferase mRNA spike-in (**Fig. 4G, and Supplementary Fig. S6A**). Additional proteins identified included nuclear speckle components such as SPOP and SRFS3, as well as nucleolar factors including the NOL8-URB1-URB2 complex, which is essential for ribosomal RNA maturation (**Fig. 4H and I**).

Together, these data show that mascRNA engages with a diverse set of tRNA-recognizing enzymes, aminoacyl-tRNA synthetases, and chromatin regulatory proteins, many of which are nuclear, nucleolar, and SUMOylated. These associations reinforce the model that mascRNA is functionally embedded within nuclear regulatory networks connected to SUMOylation, with potential links to chromatin remodeling and ribosome biogenesis.

### mascRNA transfection impacts nuclear protein SUMOylation and PML body morphology

The cumulative evidence up to this point suggested that mascRNA may modulate SUMOylation in cells. To uncover potential targets of this function, we first focused on the PBAF chromatin-remodeling complex, which was among the most enriched assemblies in the mascRNA-biotin pull-downs. Given that mascRNA associates both with the SUMO E1 enzyme (SAE1/2) and with multiple PBAF subunits, we hypothesized that this interaction could influence the SUMOylation state of PBAF components. Notably, proteomic surveys have identified several PBAF subunits including PBRM1, SMARCA4, and ARID2, as SUMO-modified proteins (42–44).

To test this hypothesis, we first performed SUMO immunoblots on PBRM1 immunoprecipitates, as well as reciprocal detection of PBRM1 in SUMO1/2 IPs, targeting either endogenous PBRM1 or a GFP-PBRM1-V5 transgene. These experiments did not yield robust or reproducible SUMO-specific signals (not shown). We next expressed FLAG-tagged SUMO1, SUMO2, and SUMO3 constructs, which increase intracellular SUMO peptide availability and allow detection of SUMOylated proteins via anti-FLAG antibodies. FLAG-SUMO1 expression markedly enhanced global SUMOylation in whole-cell lysates, whereas FLAG-SUMO2 and FLAG-SUMO3 had a weaker effect (**Fig. 5A, Supplementary Fig. S6B and C**). Increased SUMOylation of KAP1, a well-established SUMO-modified transcriptional repressor (45), verified the effectiveness of this system (**Fig. 5B**). Importantly, FLAG-SUMO1 IP followed by PBRM1 immunoblotting revealed a distinct high-molecular-weight signal, migrating at higher MW than unmodified PBRM1 only in FLAG-SUMO-expressing cells, strongly supporting that PBRM1 is SUMOylated (**Fig. 5C**). However, the relatively low yield of PBRM1 recovery and inherent variability of FLAG IPs precluded precise quantification of PBRM1 SUMOylation across conditions, and the impact of mascRNA transfection. The same approach for two additional PBAF components, ARID2 and BRG1/SMARCA4 could not confirm the presence of the PTM (**Supplementary Fig. S6D and E**), and thus this line of analysis was not pursued further.

**Figure 5.**
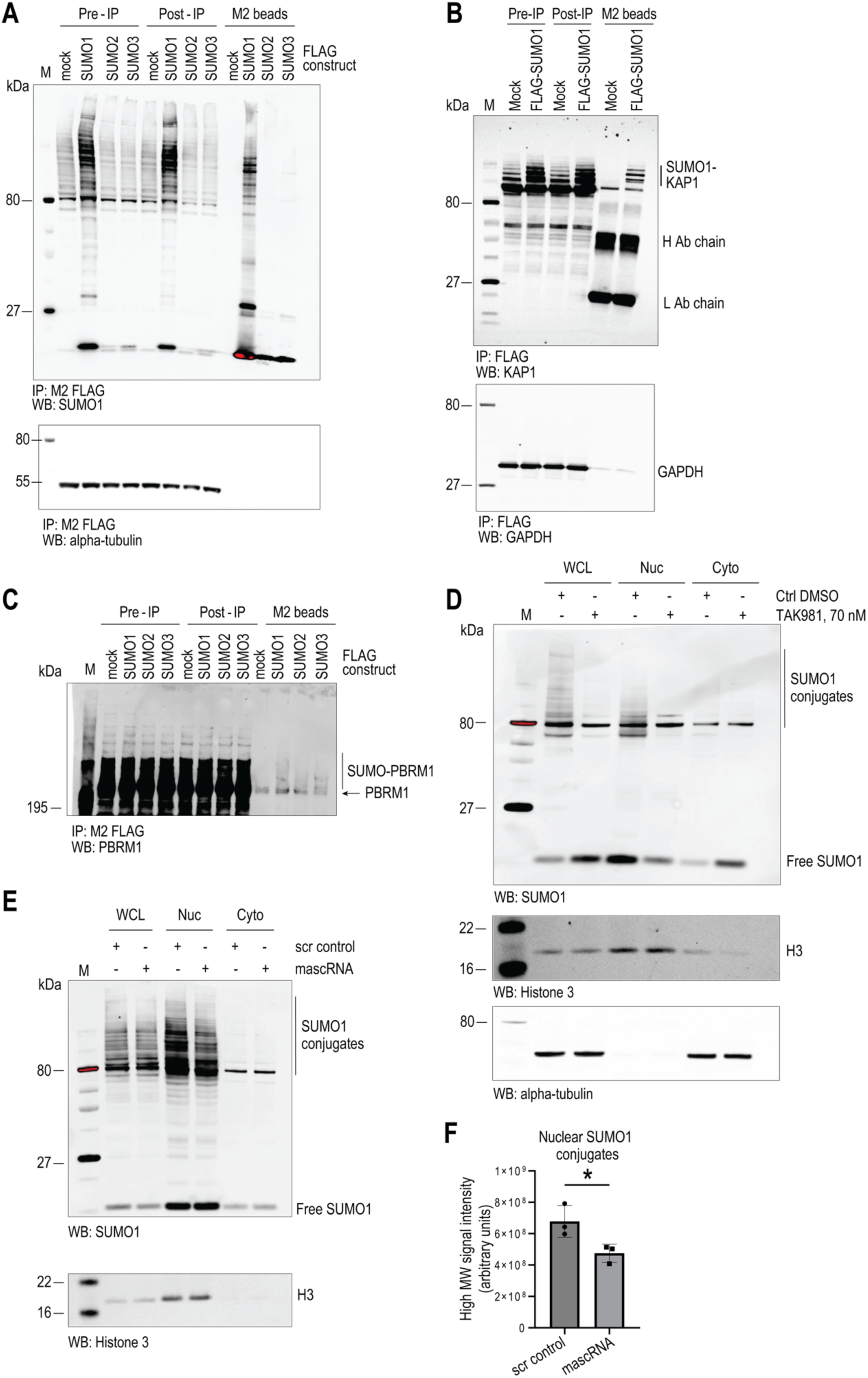
Detection of SUMOylated proteins and modulation of nuclear SUMOylation by mascRNA transfection. (**A**). Transient FLAG-SUMO1 expression for the tagging and immunocapture of SUMOylated proteins. (**B**). Immunoblot of SUMOylated KAP1 protein from FLAG-SUMO1 immunoprecipitates. (**C**). Immunoblot of SUMOylated PBRM1 protein from FLAG-SUMO1 immunoprecipitates. (**D**). Inhibition of SUMOylation in cell culture by TAK981. Cellular fractions were blotted for SUMO1. Histone3 and alpha-tubulin were used as controls of cell fractionation. WCL: whole cell lysate, Nuc: nuclear fraction, Cyto: cytoplasmic fraction. (**E**). MascRNA transfection induces reduced nuclear protein SUMOylation. Scrambled oligo (scr) was used as a control, cell fractionation as in (D). SUMO1 conjugates are highlighted. One representative replicate experiment shown, see also Supplementary Fig. S6F and G. (**F**). Quantification of SUMO1 conjugates (area highlighted in (E)) from three replicate mascRNA vs scrambled control oligo experiments. Data are shown mean ± S.D of densitometric signal at MW > 80 kDa, Student’s *t*-test, \**p*=0.040, error bars: SD.

We next turned to assessing global and subcellular SUMOylation patterns to investigate whether exogenous mascRNA affects overall SUMO conjugation. As established (46–48), most SUMO-modified proteins show nuclear partitioning (**Fig. 5D**). As a benchmark for SUMO modulation, treatment with the E1 inhibitor TAK981 produced the expected reduction of high molecular weight SUMO1 conjugates and redistribution of free SUMO1 from nucleus to cytoplasm (**Fig. 5D**). In contrast, transfection of mascRNA consistently reduced nuclear SUMO1 conjugates at high MW (> 80 kDa), without altering the predominantly nuclear localization of free SUMO1 (**Fig. 5E and F**, and **Supplementary Fig. S6F and G**), directly supporting a role for mascRNA in modulating protein SUMOylation *in cellulo*. We sought to determine differentially SUMOylated proteins in cells treated with mascRNA vs a scrambled oligo control, using a SUMO1 IP approach. Among the antibodies tested, anti-SUMO1 from Abcam (ab32058) gave a distinct protein-band pattern in Coomassie stained gels in IP experiments compared to an IgG control experiment, and proteins were efficiently eluted with urea (**Fig. 6A**). We then proceeded to analyze urea eluates from three replicate SUMO1 and IgG control IP experiments for cells treated with mascRNA vs a scrambled oligo control for a total of 12 samples (**Fig. 6B**). SUMO1 and KAP1 immunoblotting verified the specific and robust enrichment of proteins carrying the PTM in SUMO1 IP eluates (**Fig. 6C**).

**Figure 6.**
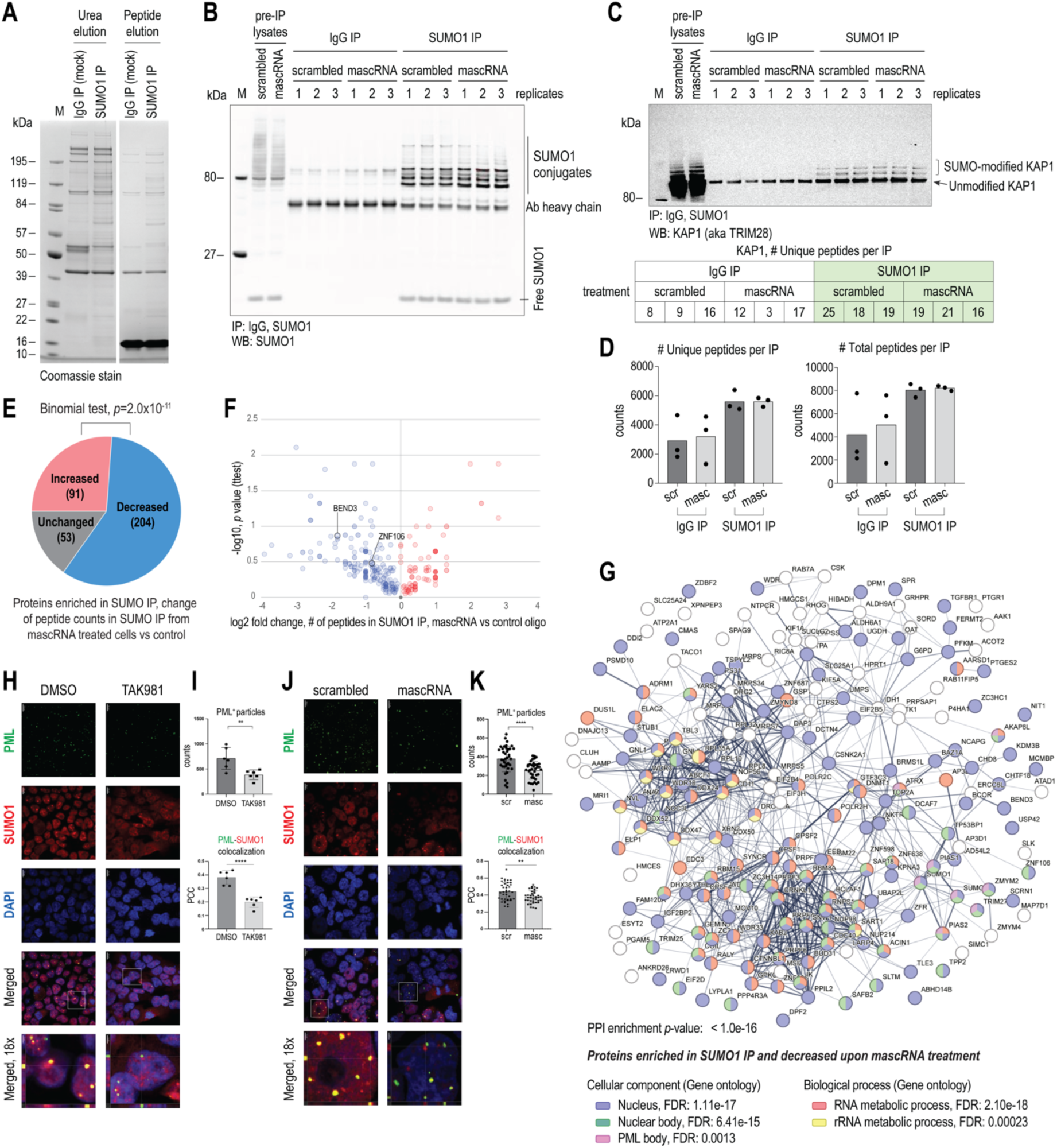
SUMO1 IP proteomics uncover *in cellulo* modulation of SUMOylation by transfected mascRNA. (**A**). SDS-PAGE analysis of Urea and SUMO1-peptide eluates of SUMO1 IP. IgG IP was used as a control. (**B**). SUMO1 immunoblot of lysates and urea eluates of SUMO1 IPs, three replicates per condition. SUMO1 conjugates are highlighted. (**C**) SUMOylated KAP1 detection in SUMO1 IPs, as a verification of the specificity of the approach. Below: table with number of KAP1 unique peptides identified in IgG and SUMO1 IPs. (**D**) Numbers of unique and total peptides identified in IP eluates of IgG and SUMO1 IP replicates. (**E**). Pie chart of proteins enriched in SUMO1 IP and how their peptide counts changed in cells treated with mascRNA vs scrambled control oligo, and binomial statistic test of decreased vs increased groups. (**F**). Volcano plot for the proteins identified in the SUMO1 IP, mascRNA vs control oligo. (**G**). STRING network analysis of the proteins that showed decreased peptide counts in the SUMO1 IP, mascRNA vs control oligo. Select statistically Gene Ontology enriched terms (FDR< 0.05) are shown; proteins are colored by term enrichment as indicated. (**H**) SUMO1 and PML immunofluorescence for co-localization in cells treated with 70 nM TAK981 SUMO E1 inhibitor, DMSO was used as a control. Scale bar, 10 μm. (**I**). Image analysis for PML positive (PML+) particles. Up: counts of PML+ particles in the two conditions. Student’s *t-*test \*\**p* <0.01, mean and ± S.D. Down: pixel-by-pixel Pearson’s Correlation Coefficient of PML and SUMO1 signal, Student’s *t-*test \*\*\*\**p* <0.0001, mean and ± S.D., *n*= 6 FOVs per condition. (**J**) SUMO1 and PML immunofluorescence for co-localization in cells treated with 10 nM mascRNA, scrambled oligo was use as a control. Scale bar, 10 μm. (**K**). Image analysis for PML positive (PML+) particles. Up: counts of PML+ particles in the two conditions. Student’s *t-*test \*\**p* <0.01, mean and ± S.D. Down: pixel-by-pixel Pearson’s Correlation Coefficient of PML and SUMO1 signal, Student’s *t-*test \*\**p* <0.01, mean and ± S.D., *n*=39 FOV 5’P-scr vs *n*=38 FOV 5’P-masc.

We identified on average ∼4900 peptides per IgG IP and ∼8300 peptides in our SUMO1 IPs, representing a total of 2518 proteins (**Fig. 6D**). To identify SUMO-modified proteins in our SUMO1 IP with high confidence, we excluded proteins identified in the SUMO1 IP represented by: a) ≥2 total peptides in any of the IgG IP samples; b) ≤5 total peptides in the SUMO1 IP replicates combined (scrambled control and mascRNA treated); c) less than 2x more peptides in the SUMO IP replicates than the IgG IP replicates, combined. These filters resulted in a list of 348 unique proteins represented by 6512 total peptides in the SUMO1 IP replicates (348 total peptides in the IgG control IPs) (**Supplementary Table S3**). This is likely an underestimate, as robustly SUMOylated and strongly enriched proteins in our SUMO1 IP will be excluded from this list if the unmodified protein shows even the slightest non-specific binding to the IgG control, as we observed for KAP1 (**Fig. 6C**).

Importantly, STRING analysis revealed that the overwhelming majority of the identified proteins had a nuclear annotation (250 out of 348, FDR 4.98×10^-31^), with significant numbers of those localized in nuclear bodies (FDR: 1.13e-22) such as nuclear speckles, the nucleolus, and the Promyelocytic Leukemia nuclear body (PML-NBs) (FDR: 6.82×10^-18^, 3.00×10^-10^, and 4.33×10^-5^ respectively) (**Supplementary Fig. S6H**).

The majority of the identified proteins (204 of 348) had on average fewer peptides in the SUMO1 IP from mascRNA-treated cells compared to the scrambled oligo control, whereas 53 proteins showed unchanged and 91 increased peptide recovery (**Fig. 6E**). This directional bias was highly significant (binomial test, *p*=2.0×10^-11^), indicating that the reduction in SUMO-associated peptides is not random but reflects a broad decrease in SUMOylation following mascRNA transfection. This result agrees with our immunoblot analysis of lysates from cells treated with mascRNA (**Fig. 5E**). Due to replicate sample variability, when individual proteins were analyzed by pairwise statistical tests across the three biological replicates per condition (two-tailed Welch’s t-test with Benjamini-Hochberg correction), no single protein passed the significance threshold adjusted for multiple testing, consistent with a modest magnitude and broad distribution of the effect across the proteome rather than large, isolated changes in a few targets (**Fig. 6F**). STRING analysis of the 204 proteins with reduced SUMOylation confirmed the significance of their localization in the nucleus (FDR: 1.11 ×10^-17^) and within nuclear membraneless organelles (FDR: 6.41×10^-15^) such as the PML bodies (FDR: 0.0013), and also showed that a significant number of these proteins are involved in RNA metabolism processes (FDR: 2.10×10^-18^) such as splicing (FDR: 1.04×10^-14^) and rRNA biogenesis (FDR: 2.3×10^-4^) (**Fig. 6G**).

We investigated the effects of the observed changes in protein SUMOylation by examining the PML body as a key nuclear body that depends on and acts as a hub of SUMOylation in the nucleus. We carried out immunofluorescence for the PML protein, the main component of these condensates, in control oligo and mascRNA treated cells, with TAK981 treatment serving as a reference condition for SUMO pathway inhibition. As expected, TAK981 led to a pronounced redistribution of SUMO1 from the nucleus to the cytoplasm (**Fig. 6H**), consistent with our immunoblot results (**Fig. 5D**). Under these conditions, we observed a significant decrease in both the number of PML-NBs and the extent of PML/SUMO1 colocalization (**Fig. 6I**). Importantly, mascRNA transfection similarly reduced PML body abundance and PML-SUMO1 colocalization relative to the scrambled control, though without changing the predominant nuclear localization of the overall SUMO1 signal (**Fig. 6J and K**). In summary, mascRNA suppresses nuclear SUMOylation and SUMO-dependent nuclear body organization without redistributing SUMO to the cytoplasm, revealing a distinct mechanism from TAK981-mediated SUMO E1 inhibition.

### mascRNA transfection impacts SUMOylation of factors involved in rRNA biogenesis

We pursued to independently validate protein SUMOylation and measure its changes upon mascRNA treatment for individual proteins, by immunoblot. For these experiments, in addition to probing the SUMO1 IP with protein specific antibodies, we used transient transfection of FLAG-SUMO1, which facilitates immunodetection of SUMO1 conjugates (**Fig. 5A**). From the list of 204 proteins showing reduced peptide numbers in SUMO1 IP from mascRNA treated cells, we focused on proteins that show nuclear body localization, and that were previously identified to be SUMOylated (42, 44). In total 13 proteins were tested by immunoblot for presence in FLAG-SUMO1 IPs with lysates from control oligo and mascRNA treated cells: BEND3, ZNF106, TOP2A, SNW1, ZC3H14, SET1, SUPT6H, IMP3, DHX36, TRIM25, CPSF7, CPSF6 and UTP18. Many of the proteins we tested either showed poor immunoblot signal or no clear enrichment in FLAG-SUMO1 IP eluates (not shown).

Two proteins, ZNF106 (aka ZFP106, an RNA-binding zinc-finger protein that interacts with GGGGCC repeat sequences and localizes to nuclear specks/nucleolus, implicated in rRNA biogenesis) (49) and BEND3 (a BEN domain containing transcription repressor that localizes in the nucleolus and is a regulator of rRNA transcription) (50) showed signals migrating above the expected MW of the unmodified proteins in SUMO1 IP immunoblots (**Fig. 7A and B**, highlighted in red). BEND3 shows a significant reduction of this high MW signal (**Fig. 7C**), while the ZNF106 signal was too diffuse to be precisely quantified. Using the FLAG-SUMO1 approach we confirmed that the high MW signals for the two proteins are SUMO1-dependent, as this signal is detected by the proteins’ respective antibodies only in FLAG-SUMO1 immunoprecipitates (**Fig. 7D and E**). To further test whether mascRNA modulates ZNF106 SUMOylation, we compared scrambled oligo and mascRNA treated cells expressing FLAG-SUMO1. Consistent with our endogenous SUMO1 IP experiments, mascRNA reduced overall SUMOylation in this setting, most clearly observed by SUMO1 immunoblotting of the FLAG IP eluates (**Fig. 7F**). Across two independent experiments, ZNF106 immunoblots showed a moderate but reproducible decrease in ZNF106-FLAG-SUMO1 conjugates in mascRNA-treated cells relative to the scrambled control (**Fig. 7G and H**). Together, these data support a model in which mascRNA dampens SUMOylation of nucleolar/nuclear-body factors linked to rRNA biogenesis and nuclear condensate organization.

**Figure 7.**
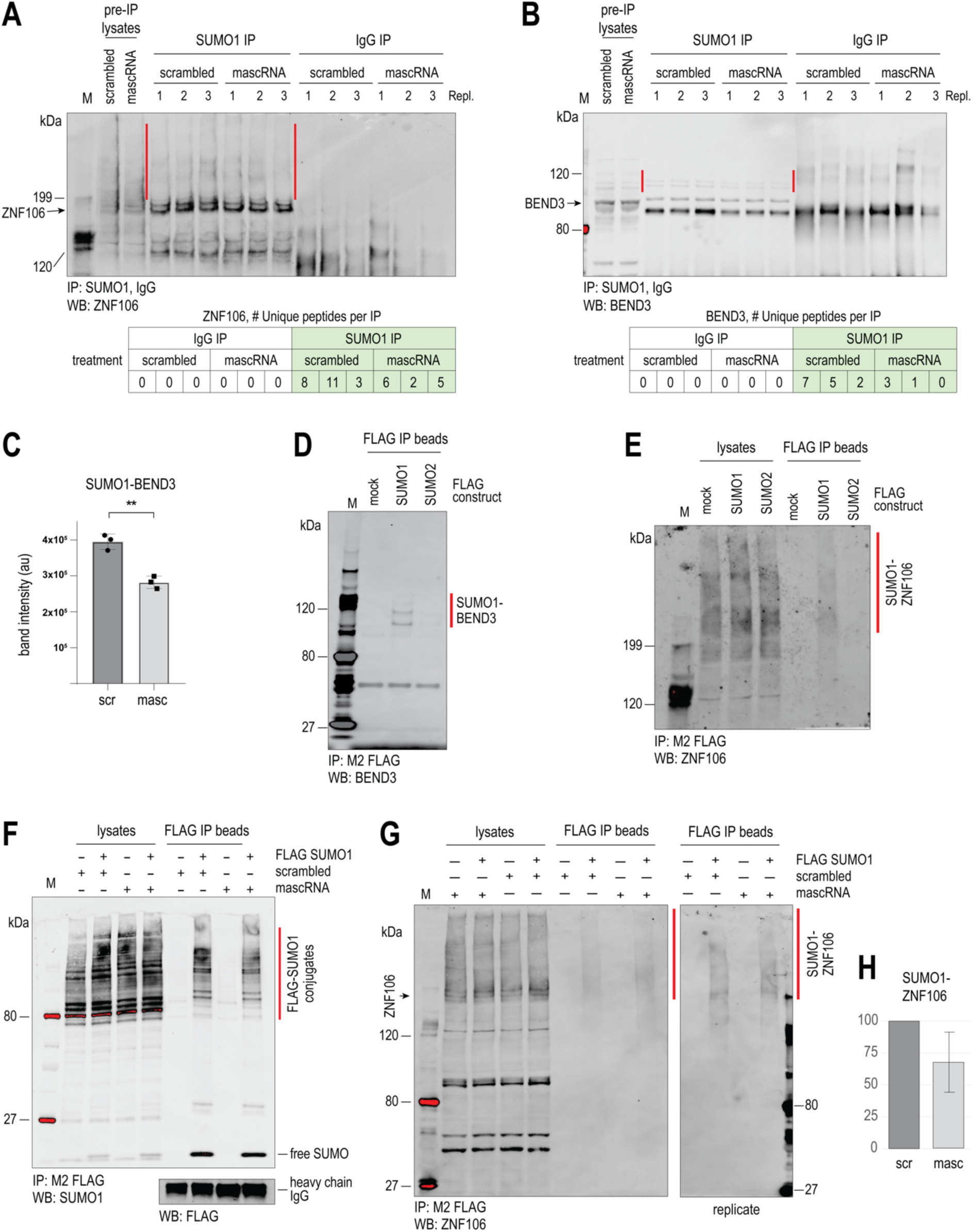
mascRNA treatment reduces protein SUMOylation. (**A**) ZNF106 immunoblot of lysates and urea eluates of SUMO1 IPs, three replicates per condition. SUMO1 conjugates are highlighted in red; below: number of unique ZNF106 peptides identified in SUMO1 IP proteomics. (**B**) BEND3 immunoblot of lysates and urea eluates of SUMO1 IPs, three replicates per condition. SUMO1 conjugates are highlighted in red; below: number of unique BEND3 peptides identified in SUMO1 IP proteomics. (**C**) Quantification of SUMO1 conjugates from panel (B). **p=0.0021, Student’s *t* test, *n*=3, error bars: SD. (**D**) Detection of SUMO1-modified BEND3 in FLAG IPs, with lysates from cells transiently expressing the FLAG constructs indicated. (**E**) Detection of SUMO1-modified ZNF106 in FLAG IPs, with lysates from cells transiently expressing the FLAG constructs indicated. (**F**) Effect of mascRNA vs scrambled oligo transfection in cells expressing FLAG-SUMO1; SUMO1 immunoblot of lysates and FLAG-IP eluates. FLAG-SUMO1 conjugates are highlighted in red. (**G**) ZNF106 immunoblot in lysates and FLAG IP eluates of FLAG-SUMO1 expressing cells, transfected with mascRNA and scrambled oligo, two replicate experiments. FLAG-SUMO1 conjugates are highlighted in red. (**H**) Quantification of SUMO1 conjugates from panel (G), normalized to scrambled oligo signal. Error bars: min-max, *n*=2.

## DISCUSSION

In this study, we define the in-solution structure and molecular interaction landscape of human mascRNA and uncover a previously unrecognized connection between this tRNA-like RNA and the SUMOylation pathway. Using complementary probing and SEC-SAXS approaches, we show that mascRNA adopts a compact cloverleaf secondary structure and an L-shaped tertiary fold in solution, consistent with but not identical to recent crystal structures of mascRNA and related processing intermediates (3, 4). In addition, we observed flexibility within the D- and anticodon-arm equivalents, and a shortened global dimension compared to canonical tRNAs. These structural features are consistent with, and help explain the selective engagement of mascRNA with precursor tRNA processing enzymes and diverse protein partners while remaining excluded from other components of the mRNA translation machinery.

Our unbiased proteomic analyses reveal that, in RNA pull-down assays, mascRNA associates with a broad set of tRNA-recognizing and tRNA-modifying enzymes, including RNase P, pseudouridine synthases, and tRNA methyltransferases, reinforcing the functional relevance of its tRNA-like architecture. In agreement with earlier work (3, 8) we also detected interactions with aminoacyl-tRNA synthetases, particularly components of the multisynthetase complex. Notably, aaRS enrichment was markedly reduced in the pull-down with a mascRNA bearing a 3’-CCA end, suggesting that this structural element modulates these interactions. Although this observation should be interpreted cautiously given the use of 3’ biotinylated RNA probes in these experiments, RNase P that also accommodates the tRNA acceptor stem in its catalytic center was recovered with comparable efficiency using both biotinylated oligos. Together, these findings support a model in which mascRNA engages aaRSs through structural features that differ from canonical tRNA recognition, permitting enzyme binding but precluding aminoacylation.

A central finding of this work is the direct association of mascRNA with the SUMO E1 activating enzyme. SAE1/2-mascRNA EMSA and RNase footprinting, along with SAE2 IP and CLIP experiments converge to demonstrate that the SAE1/2 complex, at least through its SAE2 subunit, possesses an intrinsic RNA-binding capacity and interacts with mascRNA. RNA binding by SAE1 was not directly evaluated due to the lack of a suitable antibody. These results reveal a previously unappreciated moonlighting function for this essential SUMO pathway component. Analysis of publicly available CLIP datasets further supports the existence of SAE2-RNA interactions in cells, including reads mapping to mascRNA (51). While the determinants and full scope of SAE1/2 RNA binding remain to be elucidated, these findings expand the growing concept that enzymes traditionally assigned to processes unrelated to RNA metabolism can also engage RNA.

Previous studies have reported that mascRNA levels can increase under various physiological or stress conditions, such as viral infection (10, 11). However, these conditions simultaneously trigger widespread cellular responses that complicate the interpretation of downstream effects of mascRNA function. To isolate the molecular consequences of mascRNA elevation, we transiently increased mascRNA levels by transfecting synthetic, structurally validated RNA rather than inducing endogenous expression. This approach enabled us to distinguish direct effects of mascRNA abundance from indirect consequences of stress or signaling pathways that also regulate mascRNA expression.

Under these conditions, mascRNA altered nuclear SAE2 and SUMO1 organization, reduced global nuclear SUMOylation, and reshaped SUMO-dependent nuclear bodies, including PML bodies. Importantly, this effect was mechanistically distinct from pharmacological SUMO E1 inhibition, as mascRNA reduced SUMO conjugates without redistribution of free SUMO to the cytoplasm. Proteomic analyses of SUMO1 immunoprecipitates further revealed that mascRNA transfection induced a modest reduction in SUMOylation across a significant number of nuclear proteins, particularly factors involved in RNA metabolism, chromatin regulation, and nucleolar function, consistent with a distributed regulatory role rather than selective inhibition of individual targets.

Our data indicate a functional intersection between mascRNA, SUMOylation, and nuclear body biology. Both SAE2 and mascRNA associate with nucleolar proteins (**Fig 3B**; **Fig. 4C, 4I**), and SAE2 immunofluorescence positions it in DAPI-depleted regions consistent with nucleolar localization (**Fig 3D**). Additionally, mascRNA transfection reduced SUMOylation of nucleolar and nuclear-body-localized factors implicated in rRNA biogenesis, a finding that we validated independently in FLAG-SUMO1 expressing cells (**Fig. 7**).

Although a direct interaction between mascRNA and the SUMO E1 complex is demonstrated here, the precise molecular mechanism by which mascRNA influences SUMOylation *in vivo* remains to be defined. Our primary cellular observation is a broad reduction in nuclear SUMOylation rather than selective inhibition of individual targets. Within the sensitivity of our *in vitro* assays, mascRNA did not inhibit SAE1/2 enzymatic activity or disrupt heterodimer formation, arguing against a simple competitive or catalytic blockade mechanism. Instead, interaction between mascRNA and SAE1/2 in cells may influence the spatial organization of the SUMO machinery, its engagement with partner proteins, or the selection and accessibility of substrate proteins. In this context, mascRNA may exhibit aptamer-like properties, acting as a structured RNA ligand that modulates SAE1/2 interactions within nuclear condensates, consistent with growing evidence that RNA can tune the assembly, composition, and functional output of phase-separated RNP compartments (52, 53). Together, these findings position mascRNA as an active participant in post-translational regulatory networks and open new perspectives on RNA-mediated control of protein modification and nuclear organization.

## DATA AVAILABILITY

The raw proteomics data is deposited in PRIDE with accession number PXD058139. The mascRNA SAXS model was deposited in SASBDB with accession number SASDXD9. All data generated for this paper are provided herein. Additional data may be made available upon reasonable request to the corresponding author.

## AUTHOR CONTRIBUTIONS

Supervised the study: C.S. (SAXS), J.R.T. (enzymatic probing), A.V. Designed experiments: E.L., J.R.T., C.S., A.V. Performed experiments: E.L. all figures apart from: J.R.T., C.S. Fig. 1 and Fig S1; K.D.T., A.M.H. Fig. 7D-H. Analyzed and interpreted data: E.L., J.R.T., C.S., A.V. Writing - original draft: E.L., C.S., J.R.T. Writing - Review & Editing: A.V. with input from C.S. and E.L.

## ACKNOWLEDGEMENTS

E.L. and A.V. are grateful to Professor Albert Scherpbier (UM) for his support. The authors would like to thank the team of the SWING-SAXS beamline at SOLEIL Synchrotron (Saint-Aubin, France) for beamtime allocation and support (BAG proposals No. 20181001, 20201046), Dr. Alexander Garvin and Dr. Joanna Morris (University of Birmingham, UK) for providing the WT 6xHis-FLAG-SUMO1/2/3 plasmids, Dr. Ross Tomaino (Taplin Mass Spectrometry Facility, Harvard), Dr. Fabrizio Donnarumma (LSU Mass Spectrometry Facility), Drs. David Burk and Rogerio Gomes (LSU Microscopy facility), Dr. Scott Herke (LSU Genomics Core), to Dr. Thaya Stoufflet (GeneLab, LSU Vet School), Jacqueline Paulsen and all members of the Vourekas lab for technical support and discussions.

## SUPPLEMENTARY FIGURES

**Supplementary Fig. S1.**
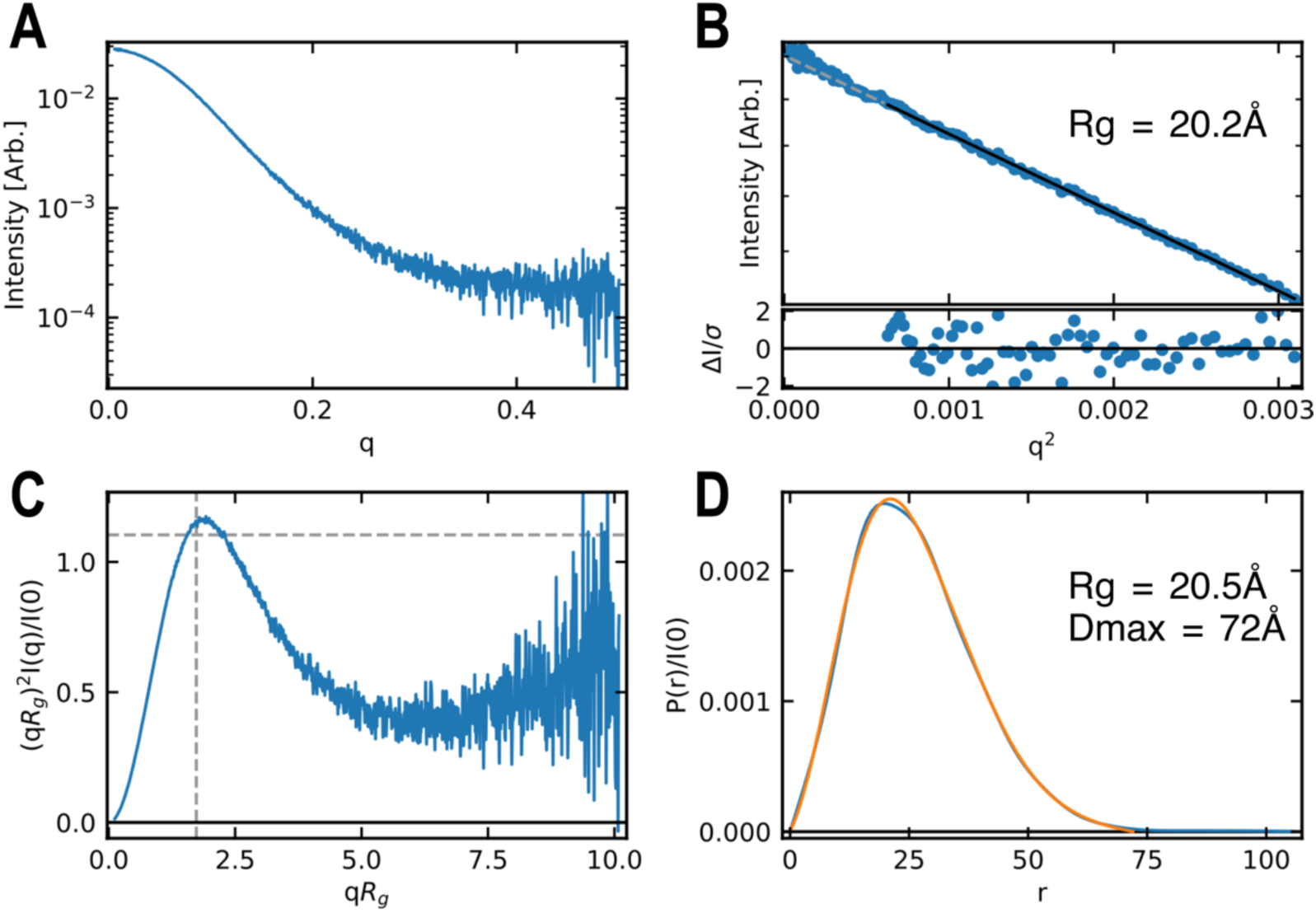
SAXS analysis of the mascRNA transcript. (**A**) Scattering curve. (**B**) Guinier plot indicating an Rg value of 20.2 Å. (**C**) Dimensionless Kratky plot showing a bell shape curve characteristic of a globular particle. (**D**) Pair-distance distribution function P(r) indicating a Rg of 20.4 Å and a Dmax of 72 Å.

**Supplementary Fig. S2.**
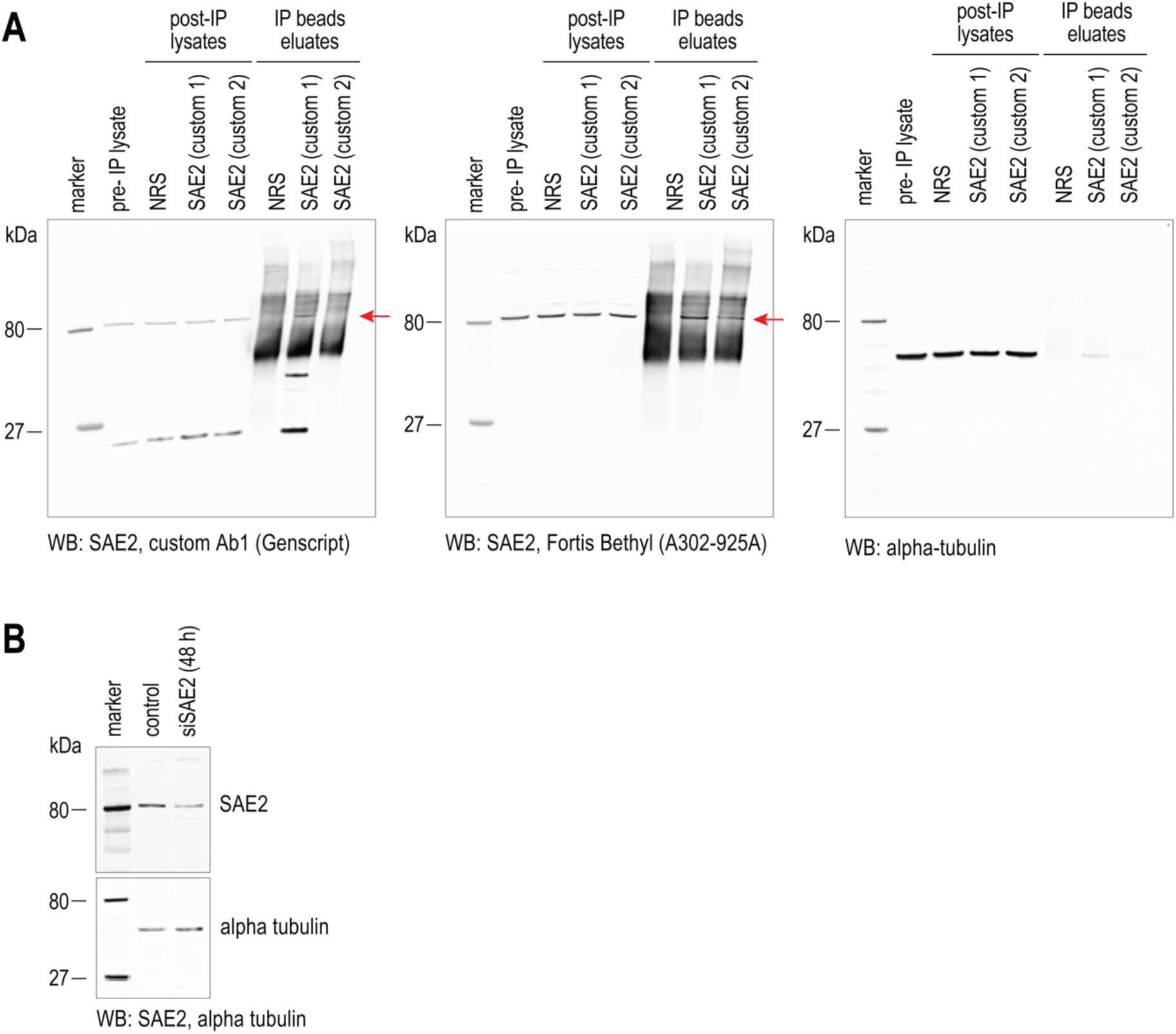
Custom anti-SAE2 Ab validation. (**A**). Validation of our custom rabbit polyclonal anti-SAE2 Ab (Ab1), with IPs and immunoblots as indicated. A red arrow points to the SAE2 band. (**B**). Validation of the SAE2 Ab1 immunoblot signal using SAE2 RNAi.

**Supplementary Fig. S3.**
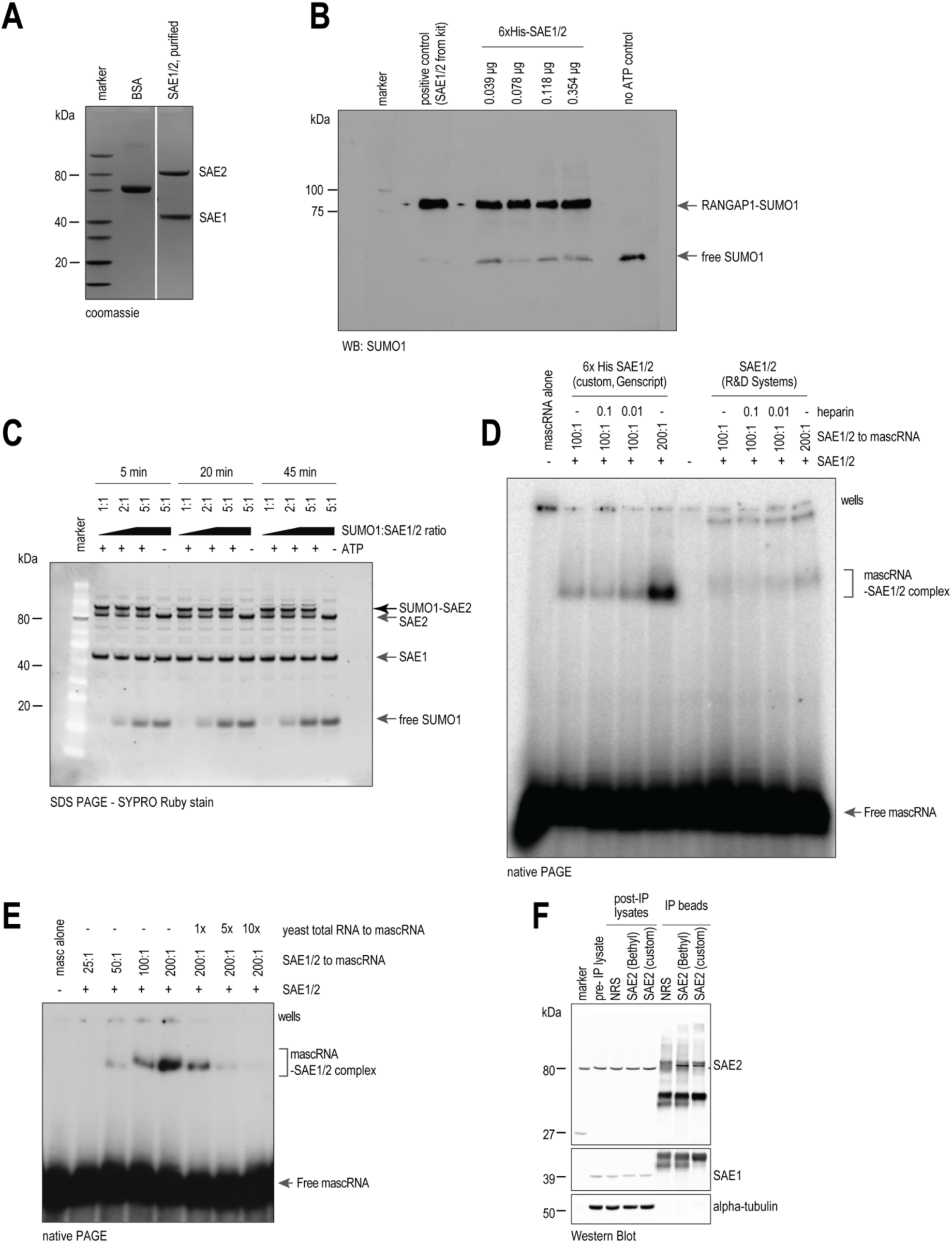
Purity and activity of recombinant SUMO E1 (SAE1/2) used for *in vitro* assays of association with synthetic mascRNA. (**A**). SDS-PAGE and Coomassie stain of purified Hist-tagged SAE1/2 complex (Genscript). (**B**). *In vitro* SUMOylation assay of purified SAE1/2 complex using RANGAP1 as a substrate, and SUMO1 immunoblot of assay products. (**C**). *In vitro* auto-SUMOylation of SAE2 by incubation of the SAE1/2 complex with SUMO1 and ATP, and SDS-PAGE analysis as indicated. (**D**). EMSA assays using SAE1/2 complex from two sources (N-terminal His-tag by Genscript, non-His-tagged from R&D) and mascRNA, as indicated. Heparin was used as indicated as non-specific polyanion competitor in some reactions. mascRNA-SAE1/2 complexes are highlighted. An additional higher MW complex was detected only with the R&D SAE1/2 preparation. The positions of the gel wells is indicated. (**E**). EMSA using SAE1/2 and mascRNA, in the presence of increasing amounts of yeast total RNA. Mass of yeast total RNA relative to mascRNA is indicated. (**F**). SAE1 and SAE2 immunoblots of lysates and IP eluates from SAE2 CLIP experiments, related to main Fig. 2F.

**Supplementary Fig. S4.**
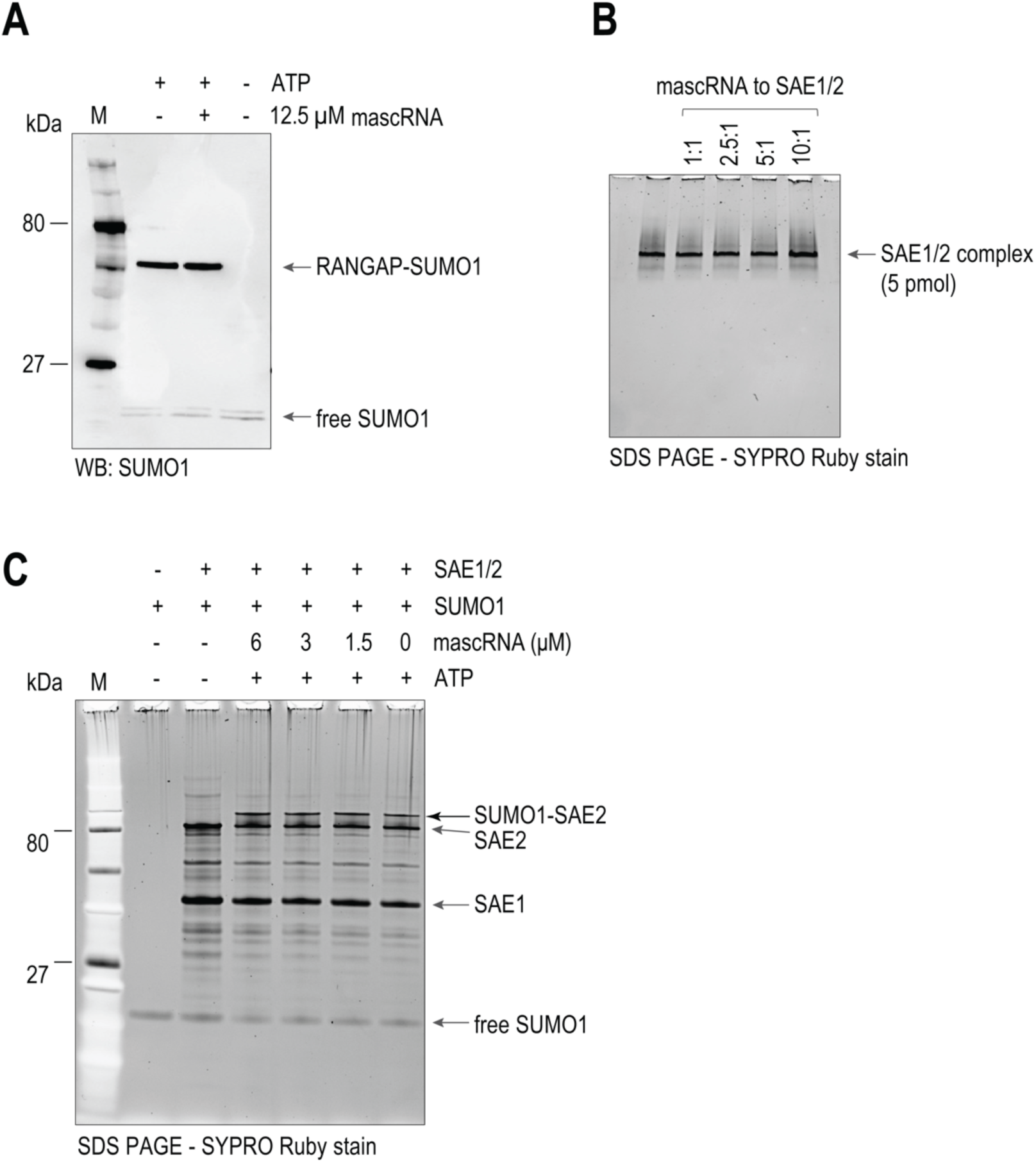
SAE1/2 in vitro activity and assembly in the presence of mascRNA. (**A**). In vitro SUMOylation assay of purified SAE1/2 complex using RANGAP1 as a substrate in the presence of mascRNA as indicated, and SUMO1 immunoblot of assay products. (**B**). Native gel analysis of the SAE1/2 complex in the absence and presence of increasing amounts of mascRNA. SAE1/2 complex as one main band, and no dissociation of the two SAE subunits in the presence of mascRNA was observed. (**C**). *In vitro* auto-SUMOylation of SAE2 by incubation of the SAE1/2 complex with SUMO1 and ATP, in the absence or presence of mascRNA, and SDS-PAGE analysis as indicated. No inhibition of the SAE1/2 enzymatic activity was observed.

**Supplementary Fig. S5.**
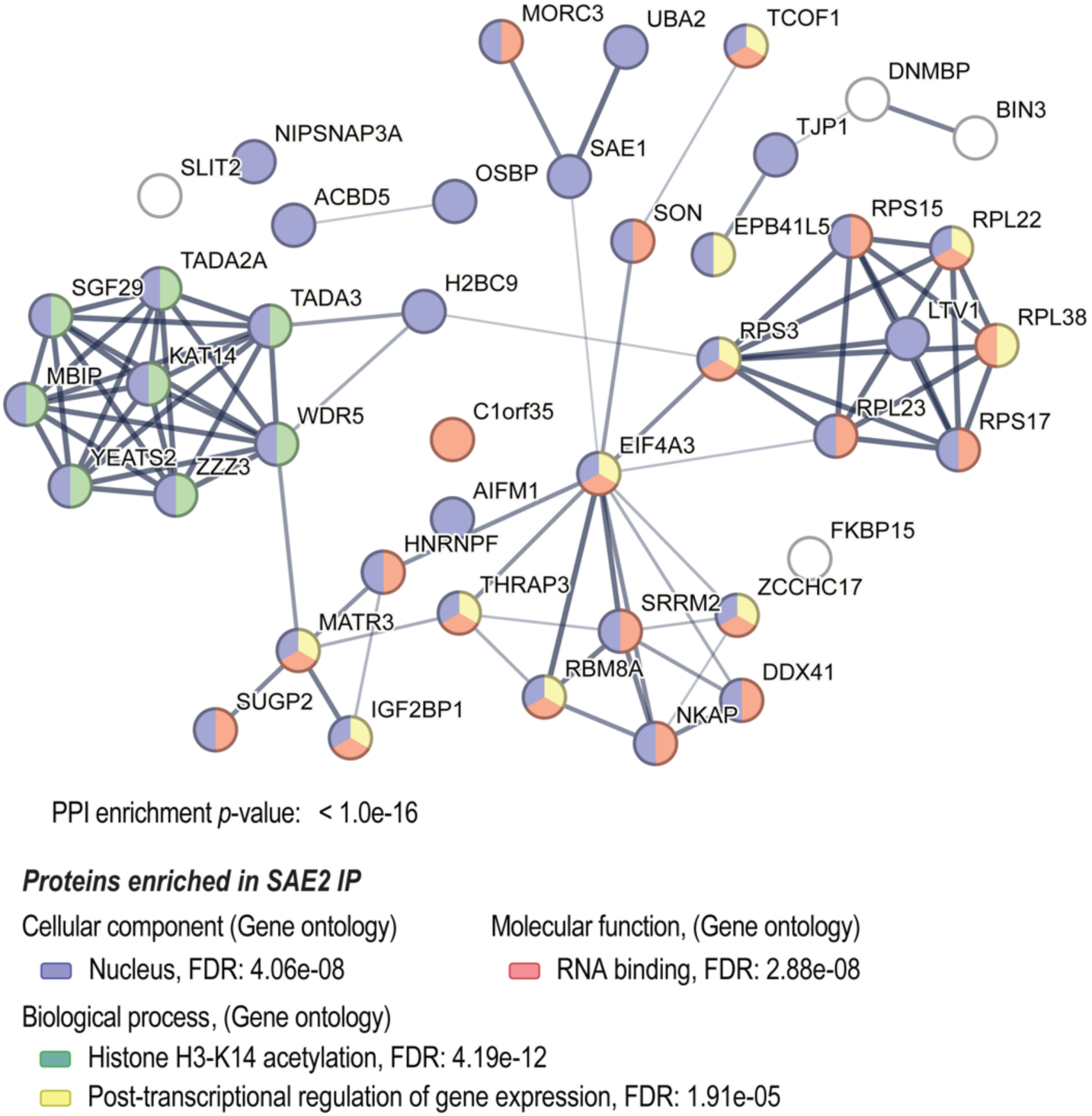
STRING network analysis of proteins enriched in SAE2 IP. Select statistically enriched Gene Ontology terms (FDR< 0.05) are shown; the proteins are colored by term enrichment as indicated.

**Supplementary Fig. S6.**
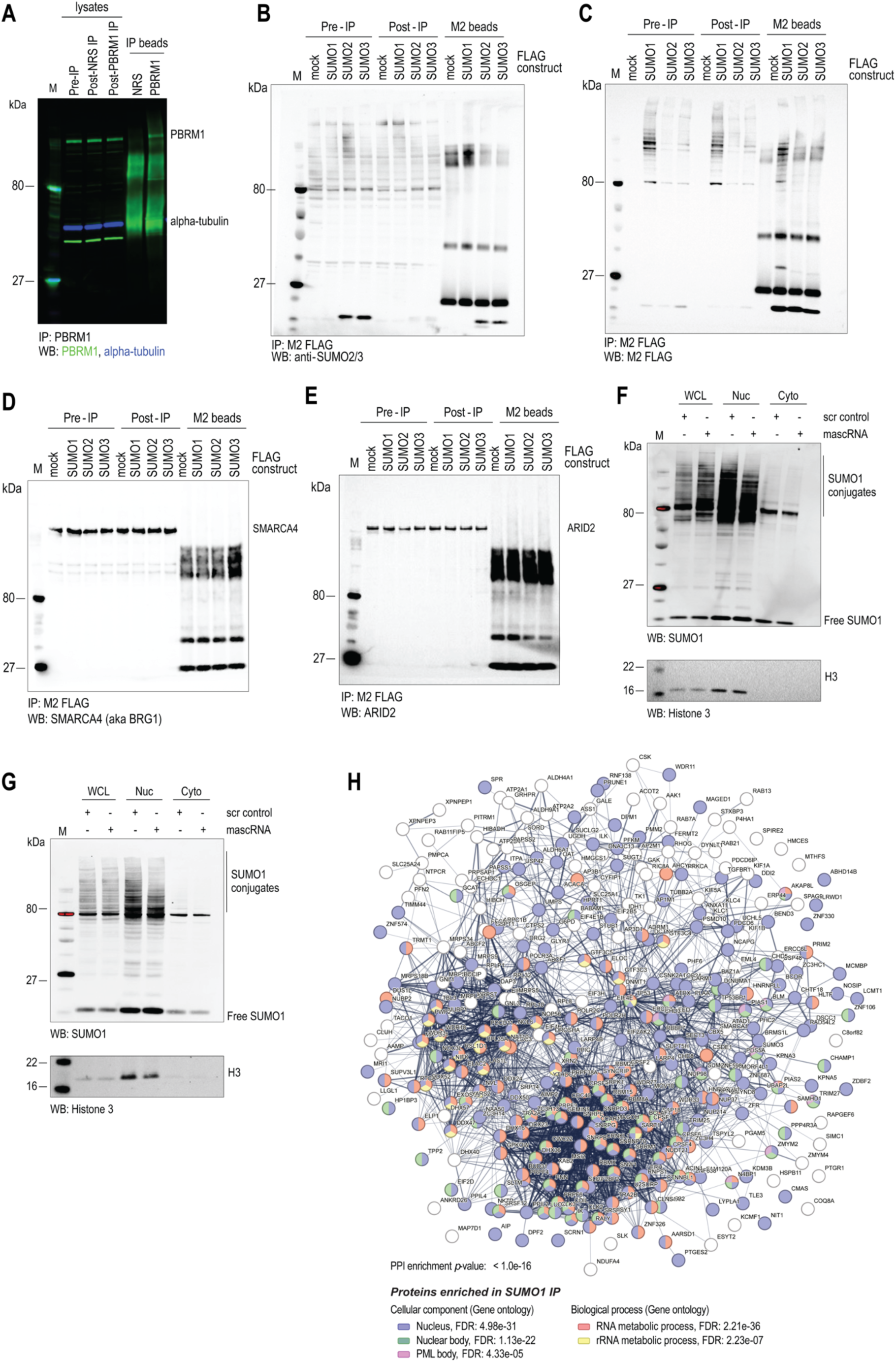
Detection of protein SUMOylation and its modulation by mascRNA transfection. (**A**). PBRM1 IP and immunoblot, related to main Fig. 4G. (**B**). Transient FLAG-SUMO2 and FLAG-SUMO3 expression for the tagging and FLAG immunocapture of SUMOylated proteins. Immunoblot with anti-SUMO2/3 Ab. (**C**). Transient FLAG-SUMO2 and FLAG-SUMO3 expression for the tagging and FLAG immunocapture of SUMOylated proteins. Immunoblot with anti-FLAG Ab (M2). (**D**). The same lysate and IP samples as shown in panels (B) and (C) were immunoblotted with anti-SMARCA4 (BRG1) Ab. Poor SMARCA4 presence in the IP eluates and no higher MW signal indicative of SUMOylation were observed. (**E**). Same as in D, immunoblot for ARID2. Poor ARID2 presence in the IP eluates and no higher MW signal indicative of SUMOylation were observed. (**F**), (**G**). MascRNA transfection induces reduced nuclear protein SUMOylation. Replicate experiments related to main Fig. 5E. Scrambled oligo (scr) was used as a control, cell fractionation as in main Fig. 5D. SUMO1 conjugates are highlighted. Histone3 was used as control of cell fractionation. WCL: whole cell lysate, Nuc: nuclear fraction, Cyto: cytoplasmic fraction. Quantifications are shown in main Fig. 5F. (**H**). STRING network analysis of proteins enriched in SUMO1 IP, from cells transfected with mascRNA or scrambled control oligo. Select statistically enriched Gene Ontology terms (FDR< 0.05) are shown; the proteins are colored by term enrichment as indicated.

## SUPPLEMENTARY TABLES

**Supplementary Table S1.** List of SAE2 (UBA2) IP mass-spectrometry identified proteins, using a commercially available SAE2 (UBA2) Antibody (Bethyl A302-925A). This list comprises proteins that were found with 0 peptides in the control NRS IP, and with 22 peptides in the SAE2 IP.

**Supplementary Table S2.** List of mass-spectrometry identified proteins using 3’ biotinylated RNA oligonucleotides: scrambled-mascRNA, mascRNA, mascRNA-CCA. This list comprises proteins with 22 peptides in any one of the two mascRNA pull-downs, and absent (0 peptides) from the scrambled control.

**Supplementary Table S3.** List of mass-spectrometry identified proteins from triplicate IPs: IgG vs. SUMO1 using lysates from scrambled mascRNA and mascRNA-transfected HEK293 cells. This list excludes proteins identified in the SUMO1 IP, and represented by: a) ≥2 total peptides in any of the IgG IP samples; b) ≤5 total peptides in the SUMO1 IP replicates combined (scrambled control and mascRNA treated); c) less than 2x more peptides in the SUMO IP replicates than the IgG IP replicates, combined.

## REFERENCES

1. Wilusz, J.E., Freier, S.M. and Spector, D.L. (2008) 3’ end processing of a long nuclear-retained noncoding RNA yields a tRNA-like cytoplasmic RNA. Cell, 135, 919–32.

2. Zhang, B., Mao, Y.S., Diermeier, S.D., Novikova, I. V., Nawrocki, E.P., Jones, T.A., Lazar, Z., Tung, C.-S., Luo, W., Eddy, S.R., et al. (2017) Identification and Characterization of a Class of MALAT1-like Genomic Loci. Cell Rep., 19, 1723–1738.

3. Skeparnias, I., Bou-Nader, C., Anastasakis, D.G., Fan, L., Wang, Y.-X., Hafner, M. and Zhang, J. (2024) Structural basis of MALAT1 RNA maturation and mascRNA biogenesis. Nat. Struct. Mol. Biol., 31, 1655–1668.

4. He, Y., Deng, J., Lin, X., Lu, Z., Wang, L., Xu, L., Zhang, Y., Wang, J. and Huang, L. (2025) Structural basis for tRNA mimicry by mascRNA and menRNA. Cell Discov., 10, 128.

5. Grozhik, A. V., Olarerin-George, A.O., Sindelar, M., Li, X., Gross, S.S. and Jaffrey, S.R. (2019) Antibody cross-reactivity accounts for widespread appearance of m1A in 5’UTRs. Nat. Commun., 10, 5126.

6. Torabi, S.-F., DeGregorio, S.J. and Steitz, J.A. (2021) tRNA-like leader-trailer interaction promotes 3′-end maturation of MALAT1. RNA, 27, 1140–1147.

7. Tripathi, V., Ellis, J.D., Shen, Z., Song, D.Y., Pan, Q., Watt, A.T., Freier, S.M., Bennett, C.F., Sharma, A., Bubulya, P.A., et al. (2010) The Nuclear-Retained Noncoding RNA MALAT1 Regulates Alternative Splicing by Modulating SR Splicing Factor Phosphorylation. Mol. Cell, 39, 925–938.

8. Lu, X., Huang, J., Wu, S., Zheng, Q., Liu, P., Feng, H., Su, X., Fu, H., Xi, Q. and Wang, G. (2020) The tRNA-like small noncoding RNA mascRNA promotes global protein translation. EMBO Rep., 21, e49684.

9. Xie, S.-J., Diao, L.-T., Cai, N., Zhang, L.-T., Xiang, S., Jia, C.-C., Qiu, D.-B., Liu, C., Sun, Y.-J., Lei, H., et al. (2021) mascRNA and its parent lncRNA MALAT1 promote proliferation and metastasis of hepatocellular carcinoma cells by activating ERK/MAPK signaling pathway. Cell Death Discov., 7, 110.

10. Gast, M., Schroen, B., Voigt, A., Haas, J., Kuehl, U., Lassner, D., Skurk, C., Escher, F., Wang, X., Kratzer, A., et al. (2016) Long noncoding RNA MALAT1-derived mascRNA is involved in cardiovascular innate immunity. J. Mol. Cell Biol., 8, 178–81.

11. Sun, T., Wei, C., Wang, D., Wang, X., Wang, J., Hu, Y. and Mao, X. (2021) The small RNA mascRNA differentially regulates TLR-induced proinflammatory and antiviral responses. JCI Insight, 6.

12. Gao, G., Wong, J., Zhang, J., Mao, I., Shravah, J., Wu, Y., Xiao, A., Li, X. and Luo, H. (2010) Proteasome activator REGgamma enhances coxsackieviral infection by facilitating p53 degradation. J. Virol., 84, 11056–66.

13. Wang, K., Jiang, Z., Lu, X., Zhang, Y., Yuan, X., Luo, D., Lin, Z., Zuo, Y. and Luo, Q. (2020) Cardiomyocyte-specific deletion of Senp2 contributes to CVB3 viral replication and inflammation. Int. Immunopharmacol., 88, 106941.

14. Goffeney, A., Hendriks, I.A., Morel, V., Loe-Mie, Y., Charon, F., Nielsen, M.L., Cossec, J.-C., Noordermeer, D., Seeler, J.-S. and Dejean, A. (2025) SUMO operates from a unique long tandem repeat to keep innate immunity in check. Nucleic Acids Res., 53.

15. Fan, Y., Li, X., Zhang, L., Zong, Z., Wang, F., Huang, J., Zeng, L., Zhang, C., Yan, H., Zhang, L., et al. (2022) SUMOylation in Viral Replication and Antiviral Defense. Advanced Science, 9.

16. El Motiam, A., Vidal, S., Seoane, R., Bouzaher, Y.H., González-Santamaría, J. and Rivas, C. (2020) SUMO and Cytoplasmic RNA Viruses: From Enemies to Best Friends. In.pp. 263–277.

17. Schimmel, J., Eifler, K., Sigurðsson, J.O., Cuijpers, S.A.G., Hendriks, I.A., Verlaan-de Vries, M., Kelstrup, C.D., Francavilla, C., Medema, R.H., Olsen, J.V., et al. (2014) Uncovering SUMOylation Dynamics during Cell-Cycle Progression Reveals FoxM1 as a Key Mitotic SUMO Target Protein. Mol. Cell, 53, 1053–1066.

18. Li, K., Xia, Y., He, J., Wang, J., Li, J., Ye, M. and Jin, X. (2023) The SUMOylation and ubiquitination crosstalk in cancer. J. Cancer Res. Clin. Oncol., 149, 16123–16146.

19. Lois, L.M. and Lima, C.D. (2005) Structures of the SUMO E1 provide mechanistic insights into SUMO activation and E2 recruitment to E1. EMBO J., 24, 439–51.

20. Geiss-Friedlander, R. and Melchior, F. (2007) Concepts in sumoylation: a decade on. Nat. Rev. Mol. Cell Biol., 8, 947–956.

21. Jin, J. (2019) Interplay between ubiquitylation and SUMOylation: Empowered by phase separation. Journal of Biological Chemistry, 294, 15235–15236.

22. Lamoliatte, F., McManus, F.P., Maarifi, G., Chelbi-Alix, M.K. and Thibault, P. (2017) Uncovering the SUMOylation and ubiquitylation crosstalk in human cells using sequential peptide immunopurification. Nat. Commun., 8, 14109.

23. Hu, Y., Li, X., Wang, D. and Mao, X. (2023) mascRNA alleviates STING-TBK1 signaling-mediated immune response through promoting ubiquitination of STING. Mol. Immunol., 154, 45–53.

24. Gast, M., Nageswaran, V., Kuss, A.W., Tzvetkova, A., Wang, X., Mochmann, L.H., Rad, P.R., Weiss, S., Simm, S., Zeller, T., et al. (2022) tRNA-like Transcripts from the NEAT1-MALAT1 Genomic Region Critically Influence Human Innate Immunity and Macrophage Functions. Cells, 11, 3970.

25. Théobald-Dietrich, A., Giegé, R. and Rudinger-Thirion, J. (2005) Evidence for the existence in mRNAs of a hairpin element responsible for ribosome dependent pyrrolysine insertion into proteins. Biochimie, 87, 813–817.

26. Thureau, A., Roblin, P. and Pérez, J. (2021) BioSAXS on the SWING beamline at Synchrotron SOLEIL. J. Appl. Crystallogr., 54, 1698–1710.

27. Fechter, P., Rudinger, J., Giegé, R. and Théobald-Dietrich, A. (1998) Ribozyme processed tRNA transcripts with unfriendly internal promoter for T7 RNA polymerase: production and activity. FEBS Lett., 436, 99–103.

28. Théobald-Dietrich, A., de Wijn, R., Rollet, K., Bluhm, A., Rudinger-Thirion, J., Paulus, C., Lorber, B., Thureau, A., Frugier, M. and Sauter, C. (2020) Structural Analysis of RNA by Small-Angle X-ray Scattering. In.pp. 189–215.

29. David, G. and Pérez, J. (2009) Combined sampler robot and high-performance liquid chromatography: a fully automated system for biological small-angle X-ray scattering experiments at the Synchrotron SOLEIL SWING beamline. J. Appl. Crystallogr., 42, 892–900.

30. Hopkins, J.B., Gillilan, R.E. and Skou, S. (2017) *BioXTAS RAW*: improvements to a free open-source program for small-angle X-ray scattering data reduction and analysis. J. Appl. Crystallogr., 50, 1545–1553.

31. Manalastas-Cantos, K., Konarev, P. V., Hajizadeh, N.R., Kikhney, A.G., Petoukhov, M. V., Molodenskiy, D.S., Panjkovich, A., Mertens, H.D.T., Gruzinov, A., Borges, C., et al. (2021) *ATSAS 3.0*: expanded functionality and new tools for small-angle scattering data analysis. J. Appl. Crystallogr., 54, 343–355.

32. Svergun, D., Barberato, C. and Koch, M.H.J. (1995) *CRYSOL* – a Program to Evaluate X-ray Solution Scattering of Biological Macromolecules from Atomic Coordinates. J. Appl. Crystallogr., 28, 768–773.

33. Jossinet, F., Ludwig, T.E. and Westhof, E. (2010) Assemble: an interactive graphical tool to analyze and build RNA architectures at the 2D and 3D levels. Bioinformatics, 26, 2057–2059.

34. Wang, W., Feng, C., Han, R., Wang, Z., Ye, L., Du, Z., Wei, H., Zhang, F., Peng, Z. and Yang, J. (2023) trRosettaRNA: automated prediction of RNA 3D structure with transformer network. Nat. Commun., 14, 7266.

35. Abramson, J., Adler, J., Dunger, J., Evans, R., Green, T., Pritzel, A., Ronneberger, O., Willmore, L., Ballard, A.J., Bambrick, J., et al. (2024) Accurate structure prediction of biomolecular interactions with AlphaFold 3. Nature, 630, 493–500.

36. Suzuki, K., Bose, P., Leong-Quong, R.Y., Fujita, D.J. and Riabowol, K. (2010) REAP: A two minute cell fractionation method. BMC Res. Notes, 3, 294.

37. Kosugi, S., Hasebe, M., Tomita, M. and Yanagawa, H. (2009) Systematic identification of cell cycle-dependent yeast nucleocytoplasmic shuttling proteins by prediction of composite motifs. Proceedings of the National Academy of Sciences, 106, 10171–10176.

38. Scott, M.S., Troshin, P. V and Barton, G.J. (2011) NoD: a Nucleolar localization sequence detector for eukaryotic and viral proteins. BMC Bioinformatics, 12, 317.

39. Erdős, G., Pajkos, M. and Dosztányi, Z. (2021) IUPred3: prediction of protein disorder enhanced with unambiguous experimental annotation and visualization of evolutionary conservation. Nucleic Acids Res., 49, W297–W303.

40. Rambo, R.P. and Tainer, J.A. (2013) Accurate assessment of mass, models and resolution by small-angle scattering. Nature, 496, 477–481.

41. Jühling, T., Duchardt-Ferner, E., Bonin, S., Wöhnert, J., Pütz, J., Florentz, C., Betat, H., Sauter, C. and Mörl, M. (2018) Small but large enough: structural properties of armless mitochondrial tRNAs from the nematode Romanomermis culicivorax. Nucleic Acids Res., 46, 9170–9180.

42. Hendriks, I.A. and Vertegaal, A.C.O. (2016) A comprehensive compilation of SUMO proteomics. Nat. Rev. Mol. Cell Biol., 17, 581–595.

43. Brechalov, A. V, Georgieva, S.G. and Soshnikova, N. V (2014) Mammalian cells contain two functionally distinct PBAF complexes incorporating different isoforms of PHF10 signature subunit. Cell Cycle, 13, 1970–1979.

44. Hendriks, I.A., Lyon, D., Young, C., Jensen, L.J., Vertegaal, A.C.O. and Nielsen, M.L. (2017) Site-specific mapping of the human SUMO proteome reveals co-modification with phosphorylation. Nat. Struct. Mol. Biol., 24, 325–336.

45. Li, X., Lin, H.H., Chen, H., Xu, X., Shih, H.-M. and Ann, D.K. (2010) SUMOylation of the transcriptional co-repressor KAP1 is regulated by the serine and threonine phosphatase PP1. Sci. Signal., 3, ra32.

46. Mahajan, R., Delphin, C., Guan, T., Gerace, L. and Melchior, F. (1997) A Small Ubiquitin-Related Polypeptide Involved in Targeting RanGAP1 to Nuclear Pore Complex Protein RanBP2. Cell, 88, 97–107.

47. Pichler, A. and Melchior, F. (2002) Ubiquitin-Related Modifier SUMO1 and Nucleocytoplasmic Transport. Traffic, 3, 381–387.

48. Moutty, M.C., Sakin, V. and Melchior, F. (2011) Importin α/β mediates nuclear import of individual SUMO E1 subunits and of the holo-enzyme. Mol. Biol. Cell, 22, 652–660.

49. Tafforeau, L., Zorbas, C., Langhendries, J.-L., Mullineux, S.-T., Stamatopoulou, V., Mullier, R., Wacheul, L. and Lafontaine, D.L.J. (2013) The Complexity of Human Ribosome Biogenesis Revealed by Systematic Nucleolar Screening of Pre-rRNA Processing Factors. Mol. Cell, 51, 539–551.

50. Khan, A., Giri, S., Wang, Y., Chakraborty, A., Ghosh, A.K., Anantharaman, A., Aggarwal, V., Sathyan, K.M., Ha, T., Prasanth, K. V., et al. (2015) BEND3 represses rDNA transcription by stabilizing a NoRC component via USP21 deubiquitinase. Proceedings of the National Academy of Sciences, 112, 8338–8343.

51. Porter, D.F., Miao, W., Yang, X., Goda, G.A., Ji, A.L., Donohue, L.K.H., Aleman, M.M., Dominguez, D. and Khavari, P.A. (2021) easyCLIP analysis of RNA-protein interactions incorporating absolute quantification. Nat. Commun., 12, 1569.

52. Maharana, S., Wang, J., Papadopoulos, D.K., Richter, D., Pozniakovsky, A., Poser, I., Bickle, M., Rizk, S., Guillén-Boixet, J., Franzmann, T.M., et al. (2018) RNA buffers the phase separation behavior of prion-like RNA binding proteins. Science (1979)., 360, 918–921.

53. Mann, J.R. and Donnelly, C.J. (2021) RNA modulates physiological and neuropathological protein phase transitions. Neuron, 109, 2663–2681.

